# A dosage-sensitive ALLO-1 network coordinates mitochondrial quality control to enable functional recovery of structurally compromised muscle

**DOI:** 10.1101/2025.10.16.682884

**Authors:** Anwesha Sarkar, Lilla Biriczová, Natalia A. Szulc, Pankaj Thapa, Remigiusz Serwa, Wojciech Pokrzywa

## Abstract

Functional recovery of muscle can precede structural repair, yet the mechanisms enabling this dissociation remain elusive. In *C. elegans*, *unc-45(m94)* mutants recover motility after heat-induced paralysis despite persisting sarcomeric disorganisation, offering a model to dissect non-contractile drivers of early recovery. Quantitative proteomics identified ALLO-1, a selective autophagy adaptor previously characterised for mediating paternal mitochondrial elimination during embryogenesis, as strongly upregulated during the recovery phase. Proteomic and genetic analyses further revealed a network of ALLO-1-associated factors, including the kinase IKKE-1 (a known ALLO-1 activator), SIP-1 (a small heat shock protein), DIM-1 (a sarcomeric tether), and CAR-1 (an ER-associated RNA-binding protein). In adult muscle, ALLO-1 restrains mitophagy to preserve mitochondrial integrity. Its depletion triggers mitochondrial fragmentation and excessive turnover, especially in the sensitised *unc-45(m94)* background. Overexpression suppresses mitophagy but reduces oxidative capacity, revealing a dosage-sensitive checkpoint. While IKKE-1 and SIP-1 promote mitochondrial stability, DIM-1 and CAR-1 facilitate turnover when ALLO-1 levels fall below threshold. Together, this regulatory module integrates proteostatic, translational, and mechanical signals to safeguard mitochondrial homeostasis and enable recovery when contractile architecture is compromised.

## INTRODUCTION

Recovery of muscle function following stress or injury is a multifaceted process that cannot be explained solely by structural repair. While the rebuilding of damaged fibres is essential for long-term regeneration, several mammalian studies have shown that strength and contractility can return before sarcomeres are fully reorganized. Early improvements have been attributed to mechanisms such as restoration of excitation-contraction coupling, redox-dependent modulation of calcium sensitivity, and changes in motor unit drive^1–5^. These findings suggest that functional recovery can rely on alternative processes that compensate for persistent myofibrillar disorganisation. Yet, the molecular basis of this phenomenon remains poorly defined, and little is known about the cellular pathways that prioritise functional restoration over structural repair.

The nematode *Caenorhabditis elegans* provides a genetically tractable system to dissect such early recovery mechanisms. Its body wall muscles exhibit conservation with vertebrate skeletal muscles, including sarcomeric organisation and chaperone-mediated myosin folding^6,7^. UNC-45 is a conserved myosin co-chaperone essential for sarcomere assembly. While invertebrates such as *C. elegans* carry a single *unc-45* gene, vertebrates encode two paralogs, with UNC-45B dedicated to striated muscle and myosin folding. In worms, the *unc-45(m94)* mutation causes paralysis and sarcomeric disorganisation at restrictive temperatures due to UNC-45 instability and impaired myosin binding, providing a tractable model to study proteostasis and recovery^8–12^. Strikingly, as we show in this study, upon return to permissive conditions these mutants rapidly regain motility despite persisting sarcomeric disorganisation. This uncoupling of structural repair from functional recovery establishes a powerful experimental model to identify the cellular processes that enable motility restoration in post-mitotic muscle.

To investigate this phenomenon, we performed quantitative proteomic profiling of *unc-45(m94)* animals during the early muscle recovery phase and identified a marked upregulation of ALLO-1. ALLO-1 was originally characterised in *C. elegans* embryos as a selective autophagy adaptor mediating paternal mitochondrial elimination via allophagy, acting upstream of the ULK complex and requiring phosphorylation by IKKE-1 (Inhibitor of NF-κB kinase epsilon subunit homolog 1), the *C. elegans* homolog of mammalian TBK1^13–15^. In contrast, our data revealed that in adult body wall muscles ALLO-1 plays a fundamentally different role: rather than promoting mitochondrial degradation, it restrains mitophagy within a dosage-sensitive regulatory network. This network integrates IKKE-1 together with SIP-1 (small heat shock protein 1)^16^, as well as two newly identified interactors: DIM-1 (disorganized muscle protein 1), an immunoglobulin superfamily protein implicated in muscle attachment and mitochondrial stability^17^, and CAR-1 (cytokinesis/apoptosis regulator 1), an endoplasmic reticulum (ER)-localised RNA-binding protein involved in mRNA stability and membrane trafficking^18,19^. Together, these findings establish a framework in which ALLO-1 coordinates mitochondrial quality control to safeguard functional recovery when structural repair is incomplete.

## RESULTS

### ALLO-1 promotes motility recovery independent of sarcomere repair

To dissect the molecular mechanisms underlying muscle recovery, we took advantage of a *C. elegans* model carrying the *unc-45(m94)* allele. At the permissive temperature (15 °C), *unc-45(m94)* animals exhibited coordinated motility and well-organised sarcomeres, whereas a shift to 25 °C for 18-20 h led to paralysis and severe disorganisation of myofilaments (Fig. 1A-B). When animals were returned to 15 °C, approximately 50 % of the population regained coordinated movement within ^10–12^ h, even though sarcomere organisation remained markedly disrupted at this stage (Fig. 1A-B). These observations establish *unc-45(m94)* as a tractable model to study the mechanisms that enable functional recovery of muscle in the absence of complete structural repair.

**Figure 1.**
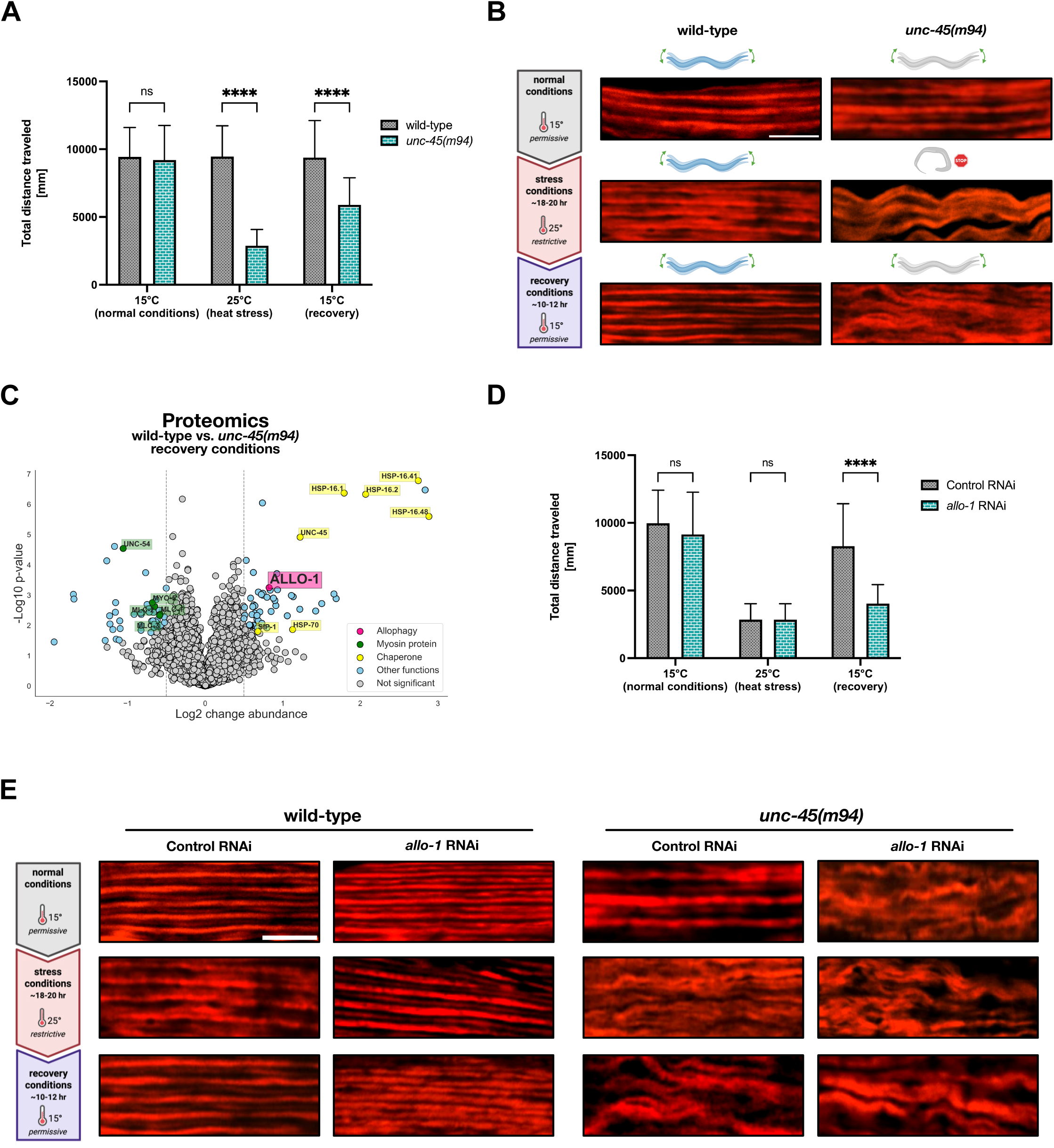
ALLO-1 promotes motility recovery independent of sarcomere repair. **(A)** Quantification of motility in young adult wild-type (N2) and *unc-45(m94)* animals. Animals were maintained at 15 °C, shifted to 25 °C, and then allowed to recover at 15 °C for 10-12 hours. Motility was measured using the WormLab system as total distance travelled over time. Data represent mean ± SEM from ≥ 3 biological replicates. Statistical analysis was performed using two-way ANOVA followed by Tukey’s multiple-comparisons test (ns - not significant; **** *P* < 0.0001). **(B)** Representative confocal images of body wall muscle stained with rhodamine-phalloidin to visualise F-actin organisation in young adult wild-type and *unc-45(m94)* animals at 15 °C, 25 °C, and after recovery at 15 °C. Scale bar, 5 µm. **(C)** Volcano plot from TMT-based quantitative proteomic analysis comparing wild-type and *unc-45(m94)* animals under recovery conditions. Statistical significance was determined using a permutation-based two-sample t-test (S₀ = 0.1, FDR < 0.05). **(D)** Quantification of motility in young adult *unc- 45(m94)* animals subjected to control (empty-vector) or *allo-1* RNAi. RNAi was initiated at the early L4 stage. Animals were maintained at 15 °C, shifted to 25 °C, and then allowed to recover at 15 °C. Motility was measured using the WormLab system as total distance travelled over time. Data represent mean ± SEM from ≥ 3 biological replicates. Statistical analysis was performed using two-way ANOVA followed by Tukey’s multiple-comparisons test (ns - not significant; **** *P* < 0.0001). **(E)** Representative confocal images of body wall muscle stained with rhodamine-phalloidin in young adult wild-type and *unc-45(m94)* animals subjected to control (empty-vector) or *allo-1* RNAi. RNAi was initiated at the early L4 stage. Animals were maintained at 15 °C, shifted to 25 °C, and then allowed to recover at 15 °C. Scale bar, 5 µm.

To identify proteins associated with the recovery phase, we performed quantitative proteomic profiling of wild-type and *unc-45(m94)* animals under three conditions: baseline at 15 °C, paralysis at 25 °C, and recovery after 10 h at 15 °C (Fig. S1A, Table S1). Gene Ontology (GO) analysis of differentially regulated proteins revealed strong enrichment of processes linked to protein folding, stress response, and actin filament organisation (Fig. S1B), reflecting coordinated activation of proteostasis and cytoskeletal maintenance pathways during recovery. Consistently, the restrictive temperature triggered a canonical stress response characterised by induction of HSP-70 and multiple HSP-16 isoforms^20,21^, which remained elevated throughout recovery (Fig. 1C, S1A). Beyond this broad chaperone upregulation, recovery was marked by the specific induction of ALLO-1, an autophagy adaptor previously characterised for its role in paternal mitochondrial elimination during *C. elegans* embryogenesis^14^ (Fig. 1C, S1A). The selective upregulation of ALLO-1 during recovery suggested a repurposed, post-developmental role in adult muscle.

To test this hypothesis, we depleted *allo-1* by RNAi initiated at the L4 stage, thereby avoiding developmental interference and focusing on the post-mitotic muscle recovery phase. Following exposure of *unc-45(m94)* animals to 25 °C and subsequent return to 15 °C, *allo-1*(RNAi) animals displayed a pronounced delay in motility recovery compared with controls (Fig. 1D). By contrast, RNAi depletion of several heat shock proteins, strongly induced during paralysis and recovery, unexpectedly accelerated recovery (Fig. S1C), suggesting that persistent activation of general stress responses may antagonise specialised mechanisms required for efficient restoration of function, whereas ALLO-1 acts within a dedicated recovery pathway.

To determine whether the requirement for ALLO-1 was linked to sarcomere integrity, we examined myofilament organisation in wild-type and *unc-45(m94)* animals following ALLO-1 knockdown. In wild-type worms, *allo-1*(RNAi) had no discernible effect on sarcomere organisation under any condition. In *unc-45(m94)* mutants, however, ALLO-1 depletion markedly aggravated myofilament disorganisation at permissive, paralytic, and recovery conditions, indicating a continuous structural role (Fig. 1E). Notably, despite these exacerbated defects, the animals remained motile at the permissive temperature (Fig. 1D), suggesting that sarcomeric disorganisation alone is insufficient to explain the observed paralysis upon heat stress and the delayed recovery. Rather, the *unc-45(m94)* background appears to reveal an additional vulnerability that is unmasked by loss of ALLO-1, pointing to non-contractile, possibly mitochondrial, mechanisms underlying the impaired recovery phase.

### ALLO-1 safeguards mitochondrial integrity, particularly in *unc-45(m94)* mutants

Given ALLO-1’s established role in paternal mitochondrial clearance during embryogenesis, we next asked whether its recovery function in adult muscle involves regulation of mitochondrial integrity. To address this, we used a TOMM-20::GFP reporter, which labels the outer mitochondrial membrane specifically in body wall muscles, enabling direct assessment of mitochondrial network morphology^22^. Mitochondria were categorised as linear, intermediate, or fragmented based on TOMM-20::GFP distribution (Fig. 2A).

**Figure 2.**
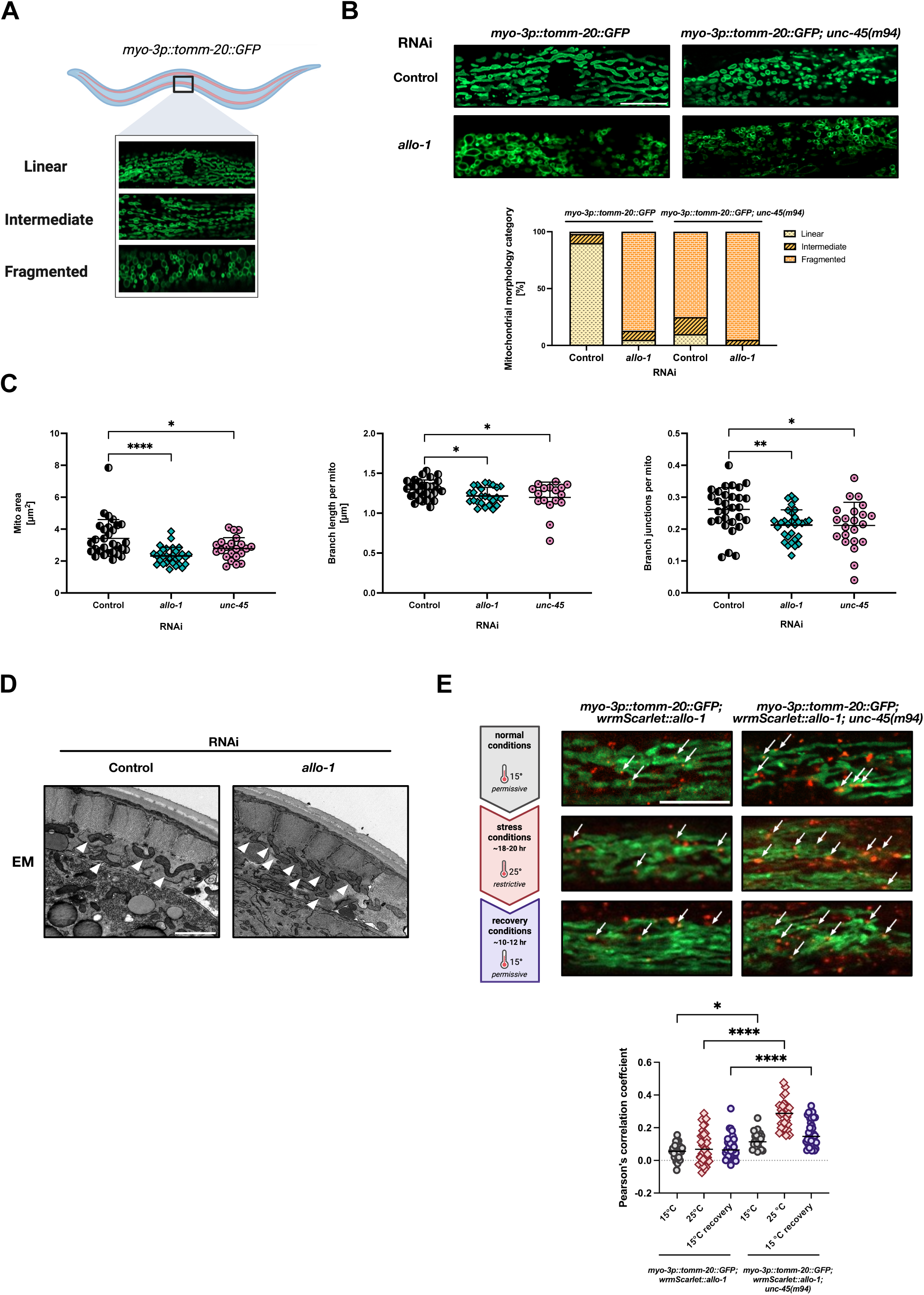
ALLO-1 safeguards mitochondrial integrity, particularly in *unc-45(m94)* mutants. **(A)** Representative confocal images of body wall muscle mitochondria in wild-type animals expressing TOMM-20::GFP. The schematic illustrates three categories of mitochondrial network morphology used for classification: linear - highly connected filamentous networks; intermediate - mixtures of punctate and elongated mitochondria; fragmented - small, rounded mitochondria with minimal branching. **(B)** Top - representative confocal images of young adult wild-type and *unc-45(m94)* animals subjected to control or *allo-1* RNAi, initiated at the early L4 stage, at 15 °C. Scale bar, 10 µm. Bottom - quantification of animals exhibiting linear, intermediate, or fragmented mitochondrial morphology (n = 20 images per condition per biological replicate; three biological replicates). **(C)** Quantitative analysis of mitochondrial morphology parameters in body wall muscle of young adult wild-type animals expressing TOMM-20::GFP after control, *allo-1*, or *unc-45* RNAi, initiated at early L4, at 20 °C. Mitochondrial area, branch length per mitochondrion, and number of branch junctions per mitochondrion were quantified. Mito - mitochondria. Data represent mean ± SEM (n ≥ 3 biological replicates). Statistical significance was assessed by one-way ANOVA with Dunnett’s multiple-comparisons test (* *P* < 0.01; ** *P* < 0.001; **** *P* < 0.0001). **(D)** Representative transmission electron microscopy (EM) images of body wall muscle mitochondria in young adult wild-type animals subjected to control or *allo-1* RNAi initiated at the early L4. White arrowheads mark individual mitochondria. Scale bar, 2 µm. **(E)** Top - Representative confocal images showing partial co-localisation of wrmScarlet::ALLO-1 (red) with TOMM-20::GFP-labelled mitochondria (green) in body wall muscle of young adult wild-type and *unc-45(m94)* animals during normal conditions of 15 °C, during stress conditions of 25 °C and during early recovery conditions of 15 °C. White arrows indicate regions of overlap. Scale bar, 10 µm. Bottom - Quantification of the degree of colocalization between GFP and wrmScarlet was performed using Pearson’s correlation coefficient. Dot plots represent data from 27-36 regions of interest (ROIs; individual cells). Statistical analysis was conducted using one-way ANOVA followed by Tukey’s multiple comparison test. (ns - not significant, *p* < 0.05, \*\*\*\**p* < 0.0001)

In wild-type animals, mitochondria were largely linear, whereas ALLO-1 knockdown induced a clear shift toward fragmented networks (Fig. 2B). A comparable, though slightly milder, fragmentation phenotype was observed upon RNAi-mediated depletion of UNC-45, suggesting that both proteins contribute to the maintenance of mitochondrial morphology under basal conditions. This cumulative effect was also apparent in *unc-45(m94)* mutants, which displayed elevated mitochondrial fragmentation already at the permissive temperature, further exacerbated by loss of ALLO-1 (Fig. 2B). To quantify these morphological changes, we performed morphometric analysis as previously described^23,24^, confirming that ALLO-1 depletion reduced mitochondrial area, branch length, and junction density (Fig. 2C). UNC-45 knockdown (RNAi) produced similar, albeit slightly less pronounced, reductions in these parameters. Transmission electron microscopy further validated this ultrastructural shift, revealing smaller, rounded mitochondria with reduced cristae complexity in *allo-1*(RNAi) animals (Fig. 2D). These data collectively indicate that both UNC-45 and ALLO-1 contribute to preserving mitochondrial architecture, and that the *unc-45(m94)* mutation sensitises mitochondria to structural disruption upon ALLO-1 loss

To further explore how ALLO-1 associates with mitochondria, we generated a CRISPR-engineered *C. elegans* strain producing endogenously tagged wrmScarlet::ALLO-1 fusion protein. Upon *allo-1* RNAi depletion, we observed a marked reduction in fluorescence, validating both the specificity of the reporter and the efficacy of the RNAi construct (Fig. S2A). Confocal imaging revealed ALLO-1 expression in body-wall muscle, where the signal co-localised with TOMM-20::GFP-labelled mitochondria in *unc-45(m94)* animals during paralysis and early recovery (Fig. 2E). This co-localisation was markedly stronger in the mutant compared to wild-type, as quantified by an increased Pearson’s correlation coefficient (Fig. 2E), indicating enhanced mitochondrial association of ALLO-1 under proteostatic stress conditions.

To test whether *allo-1* depletion broadly impairs proteostasis, we used the polyQ40::YFP aggregation reporter expressed in the body wall muscles^25^. We quantified both the number and size of aggregates and found no increase upon *allo-1* knockdown (Fig. S2B), indicating that ALLO-1 requirement during muscle recovery does not reflect a global collapse of protein folding or clearance pathways, but instead points to a specific function in mitochondrial quality control. Next, since germline signalling can also influence somatic proteostasis, we assessed fertility. In *C. elegans*, GLP-1/Notch signalling maintains the germline stem cell pool and regulates systemic stress responses^26–28^. In our assays, *glp-1(RNAi)* reduced brood size, confirming germline loss. *allo-1(RNAi)* did not affect fertility (Fig. S2C), ruling out germline-mediated effects and indicating a muscle-specific role for ALLO-1.

### Divergent ALLO-1 interactors modulate mitochondrial integrity during recovery

To gain mechanistic insights into how ALLO-1 promotes muscle recovery, we mapped its protein interactome. Immunoprecipitation of endogenous wrmScarlet::ALLO-1 followed by mass spectrometry identified both established and previously uncharacterised partners (Fig. 3A-B, S3A). Among known interactors, we confirmed IKKE-1, the *C. elegans* homolog of mammalian TBK1, which phosphorylates ALLO-1 and is required for allophagy during paternal mitochondrial clearance^13,13^. We also detected SIP-1, a small heat shock protein best characterised in the germline, where it forms pH-sensitive oligomers to counteract proteotoxic stress^16^. SIP-1 had been reported as a candidate ALLO-1 interactor in an earlier proteomic screen^16^, and in our proteomics dataset it was selectively upregulated during muscle recovery (Table S2). We validated the interaction between SIP-1 and ALLO-1 by Western blotting following immunoprecipitation, taking advantage of the availability of SIP-1-specific antibodies^16^ (Fig. S3A).

**Figure 3.**
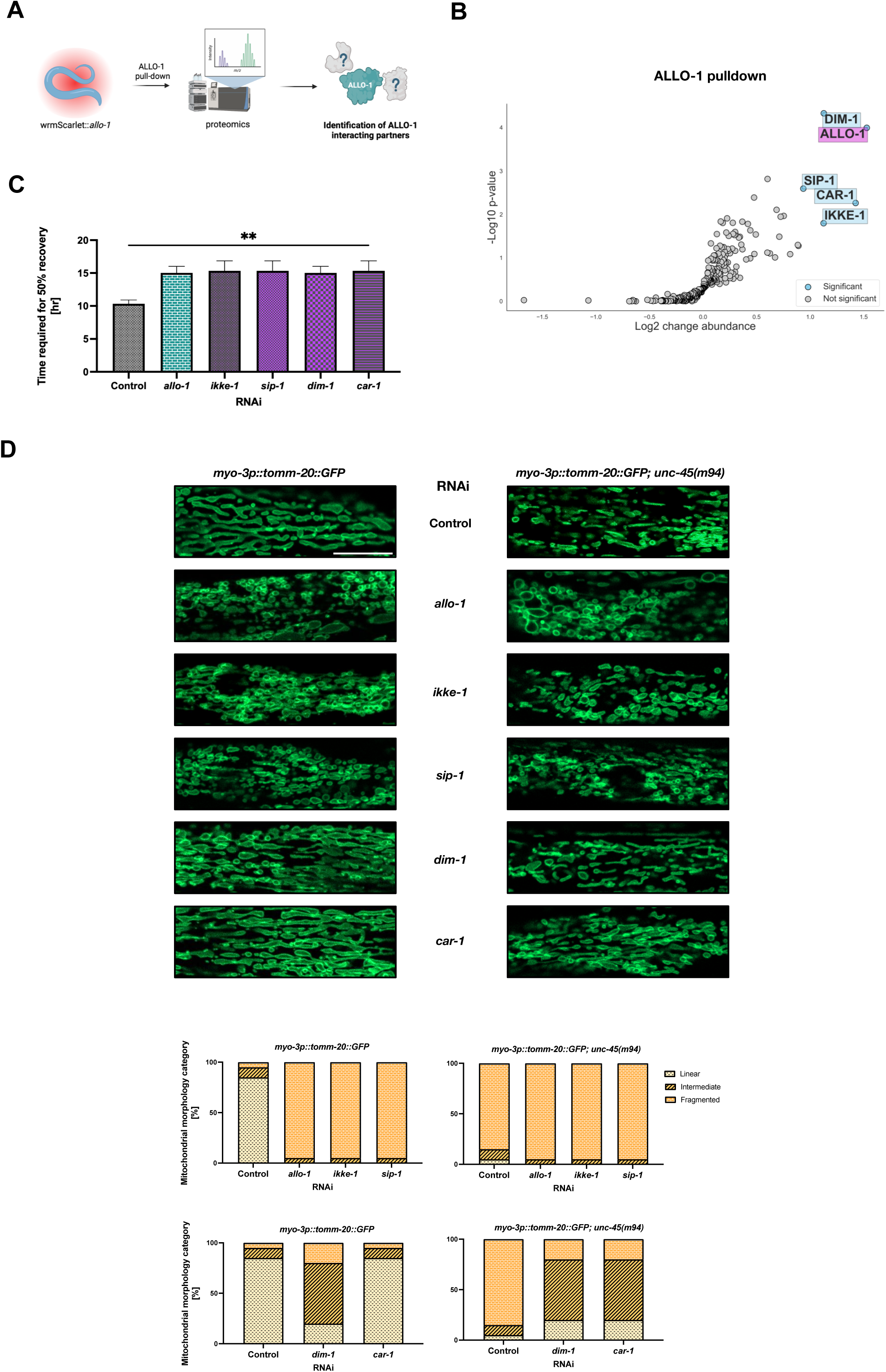
Divergent ALLO-1 interactors modulate mitochondrial integrity during recovery. **(A)** Schematic of the immunoprecipitation mass spectrometry workflow used to identify ALLO-1-interacting proteins. Lysates from young adult animals producing endogenously tagged wrmScarlet::ALLO-1 were incubated with anti-RFP beads that bind wrmScarlet. Bound proteins were subjected to on-bead digestion with trypsin, the resulting peptides were TMT-labelled and analysed by LC-MS/MS. **(B)** Volcano plot of proteins identified in wrmScarlet::ALLO-1 pulldown compared to control lysates. Statistical significance was assessed using a permutation-based *t*-test (one-sided S₀ = 0.5; FDR = 0.01). Selected interactors (*ikke-1*, *sip-1*, *dim-1*, *car-1*) are indicated. **(C)** Quantification of the time required for ∼50 % of the *unc-45(m94)* population to recover motility after down-shift from 25 °C to 15 °C under control, *allo-1*, *ikke-1*, *sip-1*, *dim-1*, or *car-1* RNAi. RNAi was initiated at the early L4 stage. Data represent mean ± SEM from ≥ 3 biological replicates. Statistical significance was determined by one-way ANOVA with Dunnett’s multiple-comparisons test (** *P* < 0.01). **(D)**Top - representative confocal images of body wall muscle mitochondria in young adult wild-type and *unc-45(m94)* animals expressing *myo-3p::tomm-20::GFP* following control or gene-specific RNAi (*allo-1*, *ikke-1*, *sip-1*, *dim-1*, *car-1*), initiated at the early L4 stage, at 15 °C. Scale bar, 10 µm. Bottom - quantification of animals exhibiting linear, intermediate, or fragmented mitochondrial morphology (n = 20 images per condition per biological replicate; three biological replicates).

Immunostaining confirmed SIP-1 expression in body wall muscles (Fig. S3B), extending its role beyond the germline and supporting a model in which SIP-1 cooperates with ALLO-1 to preserve mitochondrial integrity under stress.

In addition, we identified two previously unrecognised ALLO-1 interactors, representing mechanistically distinct modules of mitochondrial regulation, DIM-1 and CAR-1 (Fig. 3B). DIM-1 is an immunoglobulin superfamily protein, containing three Ig-like domains, that localises near dense bodies, where it contributes to anchoring actin filaments. Loss-of-function *dim-1* mutants display subtle sarcomeric disorganisation together with mitochondrial fragmentation^17^, implicating DIM-1 as a factor that couples structural stability to organellar homeostasis. By contrast, CAR-1 is an ER-associated RNA-binding protein that localises to P granules and cytoplasmic RNA foci^19^. It regulates mRNA stability, cytokinesis, and neuronal regeneration, in part through control of mitochondrial calcium uniporter components^18^, thereby providing a direct link between RNA metabolism and mitochondrial physiology. To assess the functional contribution of ALLO-1 interactors, we depleted *ikke-1*, *sip-1*, *dim-1*, or *car-1* by RNAi initiated at the early L4 stage in *unc-45(m94)* animals and monitored recovery after temperature-induced paralysis. Loss of any of these factors significantly delayed motility restoration, closely resembling the effect of *allo-1* depletion (Fig. 3C).

We next examined mitochondrial morphology in wild-type and *unc-45(m94)* backgrounds. Even under permissive conditions, *unc-45(m94)* animals exhibited a fragmented mitochondrial network (Fig. 3D). Depletion of *allo-1*, *ikke-1*, or *sip-1* further enhanced this fragmentation in both strains. We also examined *epg-7/atg-11*, the *C. elegans* homologue of mammalian FIP200/ATG11, a scaffold protein that coordinates cargo recognition with autophagosome formation and was previously implicated in positive-feedback regulation of ALLO-1 during embryonic allophagy^13^. *epg-7*(RNAi) produced only mild mitochondrial defects compared with the pronounced phenotypes of *allo-1*, *ikke-1*, or *sip-1* depletion, indicating that EPG-7 contributes less to mitochondrial maintenance in adult muscle than these core ALLO-1-associated factors (Fig. S3C). By contrast, *dim-1* depletion caused a mild shift toward intermediate mitochondrial morphologies in wild-type animals, whereas *car-1*(RNAi) had little effect. In *unc-45(m94)* mutants, however, depletion of either *dim-1* or *car-1* partially rescued the extensive fragmentation phenotype (Fig. 3D). These results thus delineate two functional classes of ALLO-1 interactors: IKKE-1 and SIP-1, which promote mitochondrial maintenance, and DIM-1 and CAR-1, which counteract fragmentation in a context-dependent manner. Together, these factors form a regulatory network in which ALLO-1 integrates both positive and buffering inputs to preserve mitochondrial organisation during muscle recovery.

### ALLO-1 and its interactors regulate mitophagy in body wall muscles

The identification of IKKE-1, SIP-1, DIM-1, and CAR-1 as ALLO-1 interactors suggested that this network may converge on pathways controlling mitochondrial quality control. Since ALLO-1 preserves mitochondrial integrity, we next examined whether it directly regulates mitophagy in muscle. To address this, we used the mitoRosella reporter, a dual-fluorescence probe targeted to the mitochondrial matrix (Fig. 4A). MitoRosella combines a pH-sensitive GFP with a pH-stable DsRed: both fluorophores are visible in intact mitochondria, whereas upon their delivery to lysosomes during mitophagy, GFP fluorescence is quenched while DsRed persists. The GFP/DsRed fluorescence ratio thus provides a ratiometric measure of mitophagic flux^29^.

**Figure 4.**
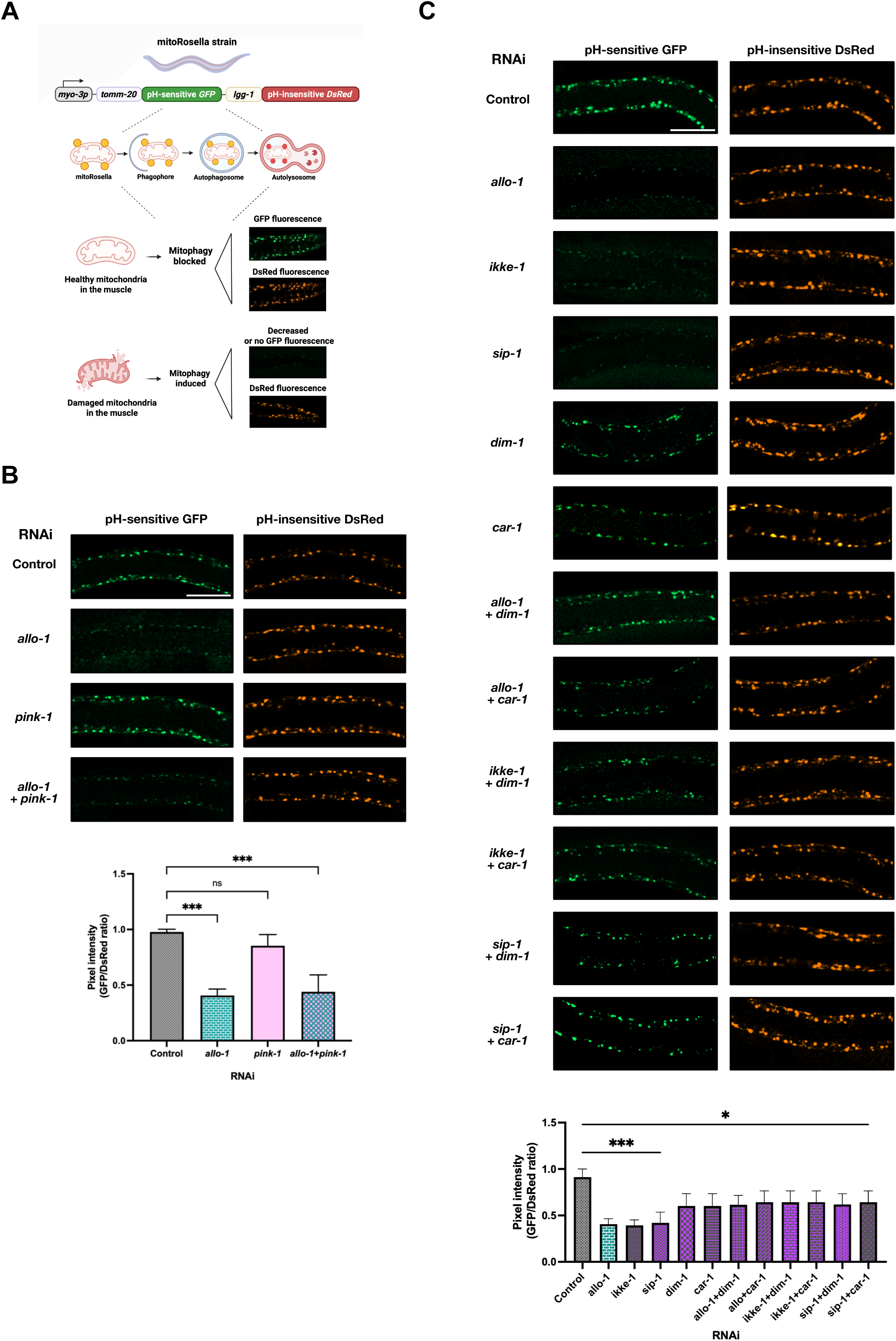
ALLO-1 and its interactors regulate mitophagy in body wall muscle. **(A)** Schematic of the mitoRosella reporter used to monitor mitophagy. **(B)** Top - representative confocal images of body wall muscle in young adult animals expressing mitoRosella reporter subjected to control, *allo-1*, *pink-1*, or combined *allo-1 + pink-1* RNAi initiated at the early L4 stage, at 20 °C. Scale bar, 50µm. Bottom - quantification of the GFP/DsRed fluorescence-intensity ratio, serving as a readout of mitophagic activity. Data represent mean ± SEM from ≥ 3 biological replicates. Statistical significance was determined using one-way ANOVA with Tukey’s multiple-comparisons test (**** *P* < 0.0001). **(C)** Top - representative confocal images of body wall muscle in young adult animals expressing mitoRosella reporter after single or combined RNAi targeting *allo-1*, *ikke-1*, *sip-1*, *dim-1*, or *car-1*, initiated at the early L4 stage, at 20 °C. Scale bar, 50µm. Bottom - quantification of the GFP/DsRed ratio. Data represent mean ± SEM from ≥ 3 biological replicates. Statistical significance was assessed by one-way ANOVA with Dunnett’s multiple-comparisons test (* *P* < 0.05; ** *P* < 0.01; **** *P* < 0.0001).

Using this system, we found that *allo-1*(RNAi) markedly increased mitophagy in body wall muscle, as shown by both microscopy and quantitative analysis of the GFP/DsRed fluorescence ratio (Fig. 4B). This finding is notable because, whereas ALLO-1 promotes mitochondrial elimination during embryogenesis^14^, in adult muscle it instead restrains mitochondrial turnover. To determine whether this induction involves the canonical PINK-1 pathway, we depleted *pink-1*, which encodes a serine/threonine kinase essential for initiating mitophagy in *C. elegans* muscle cells^30^. In wild-type animals, *pink-1*(RNAi) alone did not alter mitophagy, and it failed to suppress the elevated mitophagy caused by *allo-1*(RNAi) (Fig. 4B). These results indicate that loss of ALLO-1 triggers mitochondrial degradation through a pathway distinct from, or parallel to, the canonical PINK-1/PDR-1-dependent mitophagy.

We next asked whether ALLO-1 interactors modulate mitophagy in muscle. RNAi against *ikke-1* or *sip-1* closely phenocopied *allo-1* knockdown, producing elevated mitophagic flux as seen in microscopy images and confirmed by GFP/DsRed ratio quantification (Fig. 4C). By contrast, RNAi against *dim-1* or *car-1* consistently suppressed the mitophagy increase triggered by ALLO-1 depletion (Fig. 4C). These opposing effects parallel their influence on mitochondrial morphology (Fig. 3D), suggesting a dual regulatory scheme: IKKE-1 and SIP-1 act together with ALLO-1 to restrain mitophagy, while DIM-1 and CAR-1 function as conditional inhibitors that limit excessive mitochondrial turnover when primary regulators are compromised.

### ALLO-1 dosage determines the balance between mitochondrial turnover and function

To validate the RNAi results and exclude off-target effects, we generated a CRISPR/Cas9 *allo-1* knockout (KO) strain. Imaging of body wall muscle with the TOMM-20::GFP reporter revealed pronounced mitochondrial fragmentation in *allo-1* KO animals, closely resembling the phenotype observed upon *allo-1*(RNAi) (Fig. S4A). Additional depletion of *ikke-1*, *sip-1*, *dim-1*, or *car-1* in the *allo-1* KO background neither rescued nor further altered the severe mitochondrial phenotype (Fig. S4A), suggesting that mitochondrial organisation reaches a structural limit beyond which its interactors can no longer exert detectable effects. This implies that the regulatory capacity of the ALLO-1 network relies on the presence of residual ALLO-1 protein.

To assess the functional consequences of complete ALLO-1 loss, we quantified mitophagy using the mitoRosella reporter. *allo-1* knockout animals displayed strongly elevated mitophagic flux in body wall muscle (Fig. 5A), consistent with the results obtained by RNAi. Additional knockdown of *ikke-1* or *sip-1* did not further increase mitophagy, supporting the idea that these factors act in the same genetic pathway as ALLO-1. By contrast, depletion of *dim-1* or *car-1*, which in the RNAi context suppressed the mitophagy increase triggered by partial ALLO-1 loss (Fig. 4C), failed to reduce the elevated flux in the knockout background (Fig. 5A). These results suggest that DIM-1 and CAR-1 are required for mitophagic activation in the context of ALLO-1 depletion, and that their inhibitory effects only manifest when a minimal threshold of ALLO-1 remains. Thus, rather than directly restraining mitophagy, DIM-1 and CAR-1 appear to function as conditional effectors that mediate stress-induced mitochondrial turnover when ALLO-1 levels drop below a critical threshold, but not when ALLO-1 is entirely absent.

**Figure 5.**
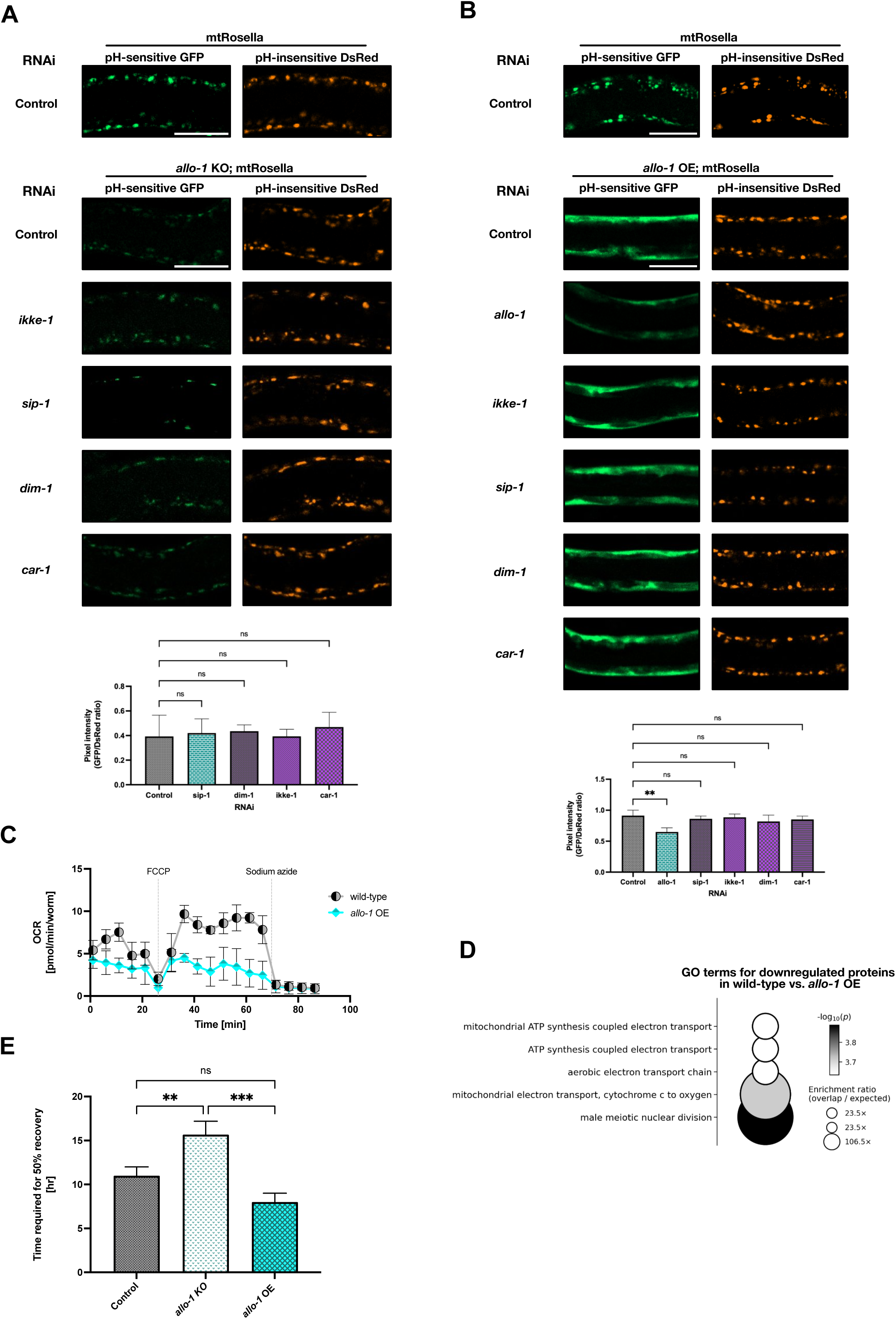
Dosage-dependent effects of ALLO-1 on mitophagy, respiration, and muscle recovery. **(A)** Top - representative confocal images of body wall muscle mitochondria in *allo-1* KO worms expressing the mitoRosella reporter, following control, *sip-1*, *dim-1*, *ikke-1* or *car-1* RNAi, initiated at the early L4 stage, at 20 °C. Scale bar, 50 µm. Bottom - quantification of the GFP/DsRed fluorescence-intensity ratios (n = 20 images per condition per biological repeat; three biological repeats). Data represent mean ± SEM and were analysed using one-way ANOVA with Dunnett’s multiple-comparisons test (ns - not significant). **(B)** Top - representative confocal images of body wall muscle mitochondria in *allo-1* OE animals expressing mitoRosella reporter, following the same RNAi treatments as in (A), at 20 °C. Scale bar, 50 µm. Bottom – quantification of the GFP/DsRed fluorescence-intensity ratios (n = 20 images per condition per biological replicate; three biological replicates). Data represent mean ± SEM. Statistical significance was determined by one-way ANOVA with Dunnett’s multiple-comparisons test (ns - not significant; ** *P* < 0.01). **(C)** Measurement of oxygen-consumption rates (OCR) in young adult wild-type and *allo-1* OE animals using a Seahorse XFp analyser, at 20 °C. Dashed lines mark compound injections (FCCP, carbonyl cyanide-4-(trifluoromethoxy)phenylhydrazone, an uncoupler of mitochondrial respiration, and sodium azide, a respiratory chain inhibitor used to block mitochondrial ATP production and assess non-mitochondrial oxygen consumption. Each point represents mean ± SEM from ≥ 8 wells (10-15 worms per well per condition). **(D)** GO biological process terms enriched among proteins downregulated in *allo-1 OE* compared with wild-type. Downregulated proteins were defined as those showing ≥ 0.3-fold decrease (S₀ = 0.1, FDR = 0.05). Enrichment analysis was performed in WebGestalt^37^ (FDR = 0.1, Benjamini- Hochberg method). Circle size denotes enrichment ratio; shading indicates statistical significance (-log₁₀ p). **(E)** Quantification of the time required for ∼ 50 % of *unc-45(m94)* animals to recover motility at 15 °C after prior exposure to 25 °C in control, *allo-1* KO, and *allo-1* OE backgrounds. Data represent mean ± SEM from ≥ 3 biological replicates. Statistical analysis used one-way ANOVA with Šidák’s multiple-comparisons test (ns - not significant; ** *P* < 0.01; *** *P* < 0.001; **** *P* < 0.0001).

To complement these loss-of-function analyses, we examined the effects of ALLO-1 overexpression (OE) in body wall muscle. We generated worms expressing the ALLO-1a isoform under the muscle-specific *myo-3* promoter (*myo-3p::FLAG-allo-1a::unc-54 3*′*UTR*), which was also used for the TOMM-20::GFP and mitoRosella reporters to ensure matched tissue specificity across assays. The ALLO-1a isoform was selected because it represents the conserved and functionally dominant variant among nematodes. During embryogenesis, ALLO- 1a localises to multiple paternal organelles, including mitochondria and membranous organelles (MOs), where it accumulates at the mitochondrial outer membrane and recruits autophagy machinery in an IKKE-1-dependent manner. In contrast, the shorter ALLO-1b isoform associates primarily with paternal mitochondria and is not conserved across nematode species^13^.

Under standard laboratory conditions (20 °C), mitochondria in ALLO-1 OE animals remained largely linear and interconnected (Fig. S4B), in contrast to the extensively fragmented networks observed in *allo-1* KO worms (Fig. S4A). Depletion of *ikke-1*, *sip-1*, *dim-1*, or *car-1* in the ALLO-1 OE background had no detectable effect on mitochondrial morphology (Fig. S4B). We next examined mitophagy using the mitoRosella reporter. In control animals, GFP⁺ signals were predominantly punctate, consistent with steady-state mitophagic turnover in body-wall muscle. By contrast, ALLO-1 OE induced a markedly diffuse and intensified GFP signal throughout the muscle cytoplasm, while the DsRed signal remained punctate (Fig. 5B). This diffuse pattern was substantially reduced upon concurrent allo-1(RNAi), confirming dose dependency, but remained unaffected by depletion of IKKE-1, SIP-1, DIM-1, or CAR-1 (Fig. 5B). Quantification of the mitoRosella GFP/DsRed ratio revealed a significant decrease in ALLO-1 OE animals, consistent with suppressed mitophagic flux. However, the emergence of a strong diffuse GFP signal suggests that this suppression may not reflect a simple block in mitophagy, but rather a more complex perturbation, such as impaired acidification, lysosomal delivery, or reporter processing. Supporting this, mitochondrial morphology visualised via TOMM-20::GFP appeared normal, without signs of fragmentation or swelling (Fig. S4B), arguing against overt mitochondrial damage or accumulation. Thus, ALLO-1 dosage critically shapes mitochondrial handling in muscle, prompting further investigation into its impact on mitochondrial function.

To evaluate how ALLO-1 OE affects mitochondrial function, we measured oxygen consumption in wild-type animals at 20 °C using Seahorse extracellular flux analysis. We focused on the overexpression model rather than the knockout, as ALLO-1 OE robustly suppresses mitophagic flux and preserves mitochondrial integrity (Fig.L5B, Fig.LS4). ALLO-1 OE worms exhibited markedly reduced basal and maximal respiration rates, indicating impaired oxidative capacity (Fig. 5C). Consistently, proteomic profiling of ALLO-1 OE animals revealed downregulation of oxidative phosphorylation and ATP synthesis factors (Fig. 5D, Table S3). Despite this respiratory limitation, ALLO-1 OE worms maintained a continuous and unfragmented mitochondrial network, with a total area comparable to that of wild type (Fig. S4B-C, Table S3). This preserved network integrity, despite reduced functional efficiency, may reflect a shift toward non-respiratory or structural roles, such as buffering proteotoxic or redox stress to maintain cellular stability during recovery. Accordingly, ALLO-1 OE animals recovered markedly faster than ALLO-1 KO worms in the *unc-45(m94)* background, and showed a slight, though not statistically significant, improvement over *unc-45(m94)* alone (Fig.L5E). The transient upregulation of ALLO-1 during the recovery phase of *unc-45(m94)* mutants (Fig.L1B) likely reflects an adaptive response that mitigates mitochondrial stress and supports contractile function before full myofibrillar repair is achieved. Together, these findings suggest that transient elevation of ALLO-1, as seen during recovery in *unc-45(m94)* mutants (Fig. 1B), represents a physiological mechanism that mitigates mitochondrial stress and supports contractile function before full myofibrillar repair is achieved.

## DISCUSSION

Our study reveals that ALLO-1, initially characterised as an adaptor mediating paternal mitochondrial elimination during embryogenesis^13,14^, adopts a strikingly different, context-dependent role in post-mitotic body-wall muscle. Rather than promoting mitophagy, ALLO-1 acts to restrain mitochondrial clearance, thereby preserving organelle integrity during recovery from proteotoxic stress. A central implication of our findings is that early functional recovery in *unc-45(m94)* mutants depends more critically on mitochondrial homeostasis than on sarcomere reassembly. Indeed, motility is regained before myofilaments are fully reassembled, indicating that intact mitochondria are the primary drivers of early recovery. Transient ALLO-1 upregulation likely serves to stabilise mitochondria, whereas its overexpression impairs oxidative capacity by suppressing mitophagy beyond an adaptive threshold. Thus, ALLO-1 acts as a dosage-sensitive checkpoint, fine-tuning mitochondrial turnover in accordance with cellular proteostatic and energetic demands.

This dosage sensitivity reflects a developmental repurposing of the ALLO-1-IKKE-1 module. In embryos, IKKE-1-dependent phosphorylation of ALLO-1 promotes mitochondrial degradation via recruitment of EPG-7/ULK14^14^. In body wall muscle, however, the same components assume a stabilising function. Depletion of either ALLO-1 or IKKE-1 results in mitochondrial fragmentation and elevated mitophagy, indicating an inversion of signalling output. IKKE-1 may maintain ALLO-1 in a phosphorylated, inhibitory state that suppresses turnover; when IKKE-1 is depleted, this restraint is lost, rendering mitochondria susceptible to degradation. Notably, *pink-1* RNAi fails to suppress the mitophagy triggered by ALLO-1 depletion, suggesting the involvement of PINK-1-independent, likely receptor-mediated or basal mitophagy pathways. The reversibility and rapid responsiveness of this regulation are consistent with known IKKE-1/ULK-driven quality-control circuits^14^. The identity of the opposing phosphatases remains to be established, but candidates such as PP2A/PPTR-2 and SCPL-1, both enriched at M-lines^31,32^, may provide additional regulatory input by antagonising IKKE-1 activity. Interestingly, SCPL-1 was found to be upregulated in the proteome of ALLO-1 OE animals (Table S3), potentially reflecting a compensatory response to ALLO-1-driven suppression of mitophagic flux or a broader rewiring of phospho-regulatory networks involved in mitochondrial quality control.

SIP-1 adds a chaperone-based layer to this regulatory network. As a small heat shock protein, SIP-1 likely buffers unfolded or oxidatively damaged proteins at mitochondrial surfaces. Its physical association with ALLO-1, combined with its upregulation in the proteomic dataset under restrictive conditions in unc-45(m94) worms and sustained expression during recovery, suggests that SIP-1 operates at the interface of protein quality control and mitochondrial maintenance. SIP-1 depletion leads to mitochondrial fragmentation and reduced autophagic flux even under physiological conditions, indicating a basal role in maintaining mitochondrial integrity. Although previously considered embryonically restricted and active under mildly acidic conditions^16^, SIP-1 is expressed in adult body-wall muscle. Local changes in pH, redox state, or ionic balance associated with sarcomere disorganisation could transiently activate SIP-1, promoting partial oligomer dissociation and increased chaperone activity. In this context, SIP-1 acts as a molecular brake on mitophagy, helping preserve mitochondrial function during transient proteostatic imbalance.

CAR-1 introduces a translational and ER-linked layer of control within the ALLO-1 network. Originally identified as an RNA-binding protein required for embryonic cytokinesis, CAR-1 regulates ER morphology and post-transcriptional gene expression^19^. In our system, CAR-1 depletion had the opposite effect to that of *allo-1*, *ikke-1*, or *sip-1* knockdown: it suppressed elevated mitophagy and mitochondrial fragmentation in *unc-45(m94)* muscle. This suggests that CAR-1 normally facilitates mitochondrial turnover, potentially by enabling local translation of factors that mediate organelle remodelling at ER-mitochondria contact sites. Comparable mechanisms have been described in neurons, where CAR-1 represses *micu-1* translation to fine-tune mitochondrial Ca^2+^ uptake after injury^18^. More broadly, mRNA storage and degradation complexes are known to associate with mitochondria and regulate their biogenesis^33^. We propose that CAR-1 functions as a post-transcriptional checkpoint, linking ER signalling to mitochondrial homeostasis. By modulating local translation, CAR-1 may facilitate mitochondrial clearance under conditions of compromised ALLO-1 activity, thereby coupling organelle turnover with translational and metabolic stress responses.

DIM-1 provides mechanical input into this regulatory network. This short immunoglobulin-like protein, localised to dense bodies and M-lines, contributes to sarcomere integrity by anchoring actin filaments to the muscle membrane^17^. Although *dim-1* loss alone causes mild lattice disorganisation, it suppresses the severe muscle disruption seen in *unc-112* mutants, suggesting a role as a mechanical stabiliser of cytoskeletal tension. In our study, DIM-1 depletion suppressed the elevated mitophagy triggered by loss of ALLO-1, IKKE-1, or SIP-1, and reduced mitochondrial fragmentation in *unc-45(m94)* mutants. This implies that DIM-1 acts as a modulator that promotes organelle turnover under conditions of reduced ALLO-1 activity. One plausible mechanism involves mechanical strain at M-lines influencing the kinase-phosphatase balance that governs ALLO-1 phosphorylation status. By providing feedback between sarcomeric strain and mitochondrial maintenance, DIM-1 may contribute to a mechanosensory checkpoint within the ALLO-1 network. Acting in parallel with CAR-1, which integrates translational cues from the ER, DIM-1 helps establish a multilayered control system that adjusts mitochondrial preservation according to the tissue’s structural and energetic status.

The reduced respiratory capacity observed in ALLO-1 overexpressing animals, despite preserved mitochondrial morphology, suggests a shift from principally bioenergetic mitochondrial states to stress-adaptive modes. Preservation of an unfragmented network may favour metabolic rewiring toward glycolytic or substrate-sparing pathways that minimise ROS burden and stabilise redox homeostasis^34,35^. Accordingly, elevated ALLO-1 during recovery may stabilise mitochondria in a semi-quiescent, structurally buffered state that sustains calcium handling and redox balance at the cost of reduced oxidative output. Both excessive and insufficient mitophagy can upset this balance and provoke functional decline. Consistent with this view, pathogenic mutations in vertebrate UNC45B result in congenital myopathies with sarcomeric disorganisation and mitochondrial loss, phenocopying aspects of *unc-45(m94)*^36^. Our discovery of a dosage-sensitive mitochondrial checkpoint in *C. elegans* thus suggests that analogous regulatory principles may underlie organelle homeostasis in vertebrate muscle, where fine-tuned mitophagy is essential to preserve proteostasis and prevent disease.

In summary, our findings identify ALLO-1 as a central, dosage-sensitive regulator of mitochondrial quality control in adult muscle. By integrating kinase, chaperone, translational, and mechanical inputs, ALLO-1 governs the balance between mitochondrial preservation and turnover, supporting tissue recovery following proteostatic challenge. This dual role, restraining mitophagy in adult muscle while promoting it in embryos, exemplifies the remarkable plasticity of mitochondrial proteostasis across developmental and physiological contexts.

## Supplementary Table Legends

Table S1. TMT-based quantitative proteomics comparing the proteomes of wild-type and *unc-45(m94)* animals during the recovery phase.

Table S2. TMT-based quantitative proteomics identifying ALLO-1 interactors.

Table S3. TMT-based quantitative proteomics comparing the proteomes of wild-type and ALLO-1 overexpression (OE) animals.

## Materials and Methods Materials availability

The mass-spectrometry proteomics datasets generated during this study have been deposited in the ProteomeXchange Consortium^38^ via the PRIDE partner repository^39^ under the accession numbers PXD069274, PXD069277 and PXD069441. The raw data were deposited in Zenodo and are available at https://doi.org/10.5281/zenodo.17371038.

### *C. elegans* culture conditions

*C. elegans* strains were maintained on nematode growth medium (NGM) plates seeded with Escherichia coli OP50 or HT115, according to standard protocols^40^. Animals were cultured at 15 °C, 20 °C, or 25 °C, depending on the strain and experimental requirements.

### *C. elegans* strains used in the study

| Genotype | Source | Identifier | Description |
| --- | --- | --- | --- |
| *C. elegans*: Bristol N2 | CGC | Strain: N2 | Wild-type worms |
| *unc-45(m94)* III. | CGC | Strain: DR94 | *unc-45(m94)* temperature-sensitive mutant |
| *wacIs14[myo-3p::tomm-20 1-50aa::attB5::mGFP::unc-54 3'UTR, unc-119(+)]*, *unc-119(ed3)* | [21] | ACH89 | TOMM-20::GFP expressed in body wall muscle |
| *myo-3p::mtRosella*; *unc-119(+)* | [28,29] | WP633 | mitoRosella reporter expressed in body wall muscle |
| *allo-1(syb8353[wrmsScarlet::allo-1])* | SunyBiotech | PHX8353 | ALLO-1 endogenously tagged with wrmsScarlet |
| *Is[myo-3p::flag-allo-1a::unc-54 3'UTR, myo-2p::gfp]* | SunyBiotech | PHX8969 | FLAG-ALLO-1a overexpression in body wall muscle with Pmyo-2::GFP marker |
| *allo-1(syb10325[allo-1(-1634bp)])* | SunyBiotech | PHX10325 | N-terminal *allo-1* knockout allele |
| *unc-45(m94) III*; *wacIs14[myo-3p::tomm-20(1-50aa)::attB5::mGFP::unc-54 3'UTR, unc-119(+)]*; *wrmsScarlet::ALLO-1* | This study | WP637 | Strain combining wrmsScarlet::ALLO-1 and *unc-45(m94)* with integrated TOMM-20::GFP reporter for mitochondrial labelling in body wall muscle |
| *wacIs14[myo-3p::tomm-20(1-50aa)::attB5::mGFP::unc-54 3'UTR, unc-119(+)]*; *wrmsScarlet::ALLO-1* | This study | WP634 | wrmsScarlet::ALLO-1 strain with integrated TOMM-20::GFP transgene for mitochondrial labelling in muscle |
| *Is[myo-3p::flag-allo-1a::unc-54 3'UTR, myo-2p::gfp]*; *wacIs14[myo-3p::tomm-20(1-50aa)::attB5::mGFP::unc-54 3'UTR, unc-119(+)]*; *unc-119(ed3) III* | This study | WP706 | Strain expressing FLAG-ALLO-1a in body wall muscle with integrated TOMM-20::GFP reporter |
| *myo-3p::mtRosella*; *Is[myo-3p::flag-allo-1a::unc-54 3'UTR, myo-2p::gfp]*; *unc-119(+)* | This study | WP705 | Body wall muscle-expressing FLAG-ALLO-1a and mtRosella reporters, with Pmyo-2::GFP as a co-injection |

|  |  |  | marker |
| --- | --- | --- | --- |
| *allo-1*(syb10325); *unc-45*(*m94*) III | This study | WP712 | Strain with *allo-1* knockout and *unc-45*(*m94*) mutation |
| *unc-45*(*m94*) III; *ls*[*myo-3p*::flag-*allo-1a*::*unc-54* 3'UTR, *myo-2p*::GFP] | This study | WP720 | Strain with FLAG-ALLO-1a extrachromosomal transgene and *unc-45*(*m94*) mutation |
| *allo-1*(syb10325); *wacIs14*[*myo-3p*::*tomm-20*(1–50 aa)::attB5::mGFP::*unc-54* 3'UTR, *unc-119*(+)];*unc-119*(*ed3*) III | This study | WP711 | Strain with *allo-1* knockout and integrated TOMM-20::GFP reporter for body wall muscle mitochondria |
| *allo-1*(syb10325); *myo-3p*::mtRosella | This study | WP714 | Strain with *allo-1* knockout and body wall muscle mtRosella reporter |

#### RNA interference

RNA interference (RNAi) was carried out using the standard feeding method, with bacterial clones sourced from the Ahringer *C. elegans* RNAi library^41^. Nematode growth medium (NGM) plates were supplemented with 1 mM IPTG (Cat. IPT001; BioShop,) and 25 µg/ml carbenicillin (Cat. 6344.2; Carl Roth GmbH & Co.,) and seeded with *Escherichia coli* HT115 (DE3) carrying the L4440 plasmid containing the target gene insert. HT115 (DE3) bacteria transformed with the empty L4440 vector served as a negative control.

#### Recovery assay for measurement of motility recovery time

Age-synchronised L4 hermaphrodites of *unc-45(m94)* were subjected to RNAi treatments as indicated in the figure legends. Animals were maintained at 15 °C and subsequently shifted to 25 °C for 18-20 h to induce paralysis. Plates were then returned to 15 °C, and motility recovery was monitored over the following ∼10 h, or until approximately 50 % of the population regained coordinated movement. *unc-45(m94)* strains carrying *allo-1* knockout or overexpression constructs were subjected to the same temperature regime and analysed in parallel to determine recovery kinetics.

#### Brood size assay

Age-synchronized L4 hermaphrodites of the wild type strain were selected and subjected to RNAi treatments targeting *glp-1, allo-1* or a control RNAi (empty vector). For the assay, two worms were placed on each 35mm NGM plate. Brood size measurements were conducted at a constant temperature of 20 °C. Worms were transferred to fresh NGM plates daily, from adulthood day 0 to adulthood day 4, marking the end of the egg-laying period. The number of eggs laid, and larvae hatched were manually counted each day throughout the reproductive stage. For each biological repeat, eggs laid, and larvae hatched from 10 worms were counted. Data are presented as the total number of eggs laid and larvae hatched per plate, aggregated across three independent biological repeats.

#### Phalloidin staining

Sarcomere organisation was analysed by staining filamentous actin (F-actin) with Phalloidin-Atto 390 (Cat. 50556; Sigma-Aldrich) following a modified protocol from^42^. Briefly, young adult *C. elegans* were washed off NGM plates seeded with *E. coli* OP50 using M9 buffer. The collected worms were snap-frozen in liquid nitrogen and dried in a SpeedVac concentrator at room temperature. Samples were permeabilised by incubation in 100 % ice-cold acetone for 5 min. After complete acetone evaporation in a fume hood, worms were incubated with Phalloidin-Atto 390 for 30 min at room temperature in the dark to label F-actin. Stained animals were washed twice with M9 buffer containing 0.05 % NP-40 (Cat. NON505; BioShop) and 5 mg/mL BSA (Cat. ALB001; BioShop) to remove unbound dye, then resuspended in a small volume of M9 and mounted on 2 % (w/v) agarose pads. Coverslips were gently applied to avoid bubble formation, and samples were imaged using a Zeiss LSM 800 inverted confocal microscope equipped with a 63× oil-immersion objective.

#### Immunostaining

Immunostaining was performed following protocols adapted from^43–46^ with minor modifications. Worms were cultured on NGM plates seeded with *E. coli* OP50 and collected at the young adult stage using M9 buffer. Samples were washed three times in M9 by centrifugation (1,000 × *g*, 2Lmin) to remove bacteria, chilled on ice, and fixed in 500 µL of ice-cold formaldehyde solution for 15-30 min at room temperature. After fixation, worms were washed three times in 1× PBS-Tween (pHL7.2) and stored at 4 °C until further processing. For permeabilization, samples were incubated overnight at 37 °C in β-mercaptoethanol (Cat.1610710; BioRad) solution with gentle mixing, followed by three washes in PBS-Tween. Collagenase (Cat. C-5138 100mg Collagenase IV Sigma) treatment was then performed at 37 °C with vigorous shaking until ∼20% of the animals appeared partially disrupted; the reaction was stopped by placing the tubes on ice. Worms were subsequently washed twice in PBS-Tween and once in antibody buffer (AbA). Primary antibody incubation was performed overnight at room temperature in AbA containing the antibody at a 1:250 dilution, with gentle agitation. The anti-SIP-1 antibody^16^ was used as the primary antibody. After incubation, worms were washed three times with AbA for 1Lh each at room temperature or overnight at 4 °C. Secondary antibody incubation was carried out in AbA at a 1:500 dilution for 1 h at room temperature in the dark, followed by three final washes in AbA. Worms were mounted on glass slides in antifade mounting medium, covered with a coverslip, and sealed with nail polish. Imaging was performed on a Zeiss LSM800 inverted confocal microscope equipped with a 63×/1.4LNA oil-immersion objective.

#### TMT-based quantitative proteomics - total protein levels

Protein extraction from *C. elegans* was performed using the Sample Preparation by Easy Extraction and Digestion (SPEED) protocol^47^. In brief, worm pellets were dissolved in concentrated trifluoroacetic acid (TFA; Cat: T6508, Sigma-Aldrich) at a pellet-to-TFA ratio of 1:2-1:4 (v/v) and incubated for 2-10 min at room temperature. Samples were then neutralised with 2 M Tris-base buffer (10× the TFA volume) and incubated at 95 °C for 5 min in the presence of 10 mM Tris(2-carboxyethyl)phosphine and 40 mM 2-chloroacetamide. Protein concentrations were estimated by turbidity measurement at 360 nm, adjusted to equal levels using dilution buffer (2 M Tris-base/TFA, 10:1 v/v), and further diluted 1:4-1:5 with water. Proteins were digested overnight at 37 °C with trypsin (protein-to-enzyme ratio 100:1), and digestion was stopped by adding TFA to a final concentration of 2%. Peptides were labelled using an on-column tandem mass tag (TMT) labelling approach^48^. Labelled samples were pooled into a single TMT multiplex, concentrated, and fractionated into 7-8 fractions using the Pierce High-pH Reversed-Phase Peptide Fractionation Kit (Cat. 84868; Thermo Fisher Scientific). Fractions were reconstituted in 0.1% TFA, 2% acetonitrile prior to LC-MS analysis. Peptide separation was performed on an Easy-Spray Acclaim PepMap column (50Lcm × 75Lµm i.d.; Cat: PN ES903; Thermo Fisher Scientific) at 55L°C using a 120-min gradient of acetonitrile in 0.1% formic acid at a flow rate of 300LnLLmin⁻¹. Mass spectrometric data were acquired on a Q Exactive HF-X Orbitrap instrument coupled to an UltiMate 3000 nano-LC via an Easy-Spray source (Thermo Fisher Scientific) operated in TMT mode. Full MS scans were acquired at 60,000 resolution (m/z 200), followed by up to 15 data-dependent MS/MS scans of precursors (charge states 2-5; isolation window 0.7Lm/z) fragmented by higher-energy collision dissociation (HCD, NCE 32). Dynamic exclusion was set to 35 s. Maximum injection times were 50 ms for MS and 96 ms for MS/MS (resolution 45,000, m/z 200). AGC targets were 3L×L10L for MS and 1L×L10L for MS/MS, with a minimum AGC of 1L×L10³. Raw data were processed in MaxQuant v1.6.17.0^49^ using the Andromeda search engine against the *C. elegans* reference proteome (Uniprot UP000001940). Reporter ion MS² quantification used a 0.003LDa mass tolerance and a minimum reporter PIF of 0.75. Modifications were set as follows: carbamidomethylation (C, fixed); oxidation (M, variable), deamidation (N/Q, variable), and N-terminal acetylation (variable). Trypsin/P was specified as the protease, allowing up to two missed cleavages. False discovery rates (FDRs) for peptides, proteins, and modification sites were set to 1%. “Match between runs” was enabled, and other parameters were kept at default settings. Quantification was based on unique and razor peptides allowing protein grouping. Reporter intensity-corrected values were imported into Perseus v1.6.10.0^50^. Reverse hits, potential contaminants, and entries identified only by site were removed. Intensities were log₂-transformed, and only protein groups with complete quantitative data across all TMT channels were retained. Normalisation was performed by median subtraction within channels. Statistical significance was determined using a two-sample t-test with a permutation-based FDR of 0.05 and S₀L=L0.1 to identify differentially expressed proteins between conditions.

#### Immunoprecipitation of ALLO-1

Immunoprecipitation of wrmScarlet::ALLO-1 was performed using RFP-Trap Magnetic Agarose (ChromoTek GmbH, Planegg-Martinsried, Germany), following the manufacturer’s protocol with minor adaptations for worm lysates. Young adult animals were collected from NGM plates using M9 buffer and washed three times by centrifugation (1,000 × *g*, 2 min) to remove bacteria. Worm pellets were frozen in liquid nitrogen and stored at −80L°C until use. Frozen pellets were homogenised in ice-cold lysis buffer (10 mM Tris-HCl pHL7.5, 150 mM NaCl, 0.5 mM EDTA, 0.5% Nonidet P40 Substitute) supplemented with 1 mM PMSF and protease inhibitor cocktail. Homogenisation was performed using a chilled motorised pestle or bead beater until a uniform lysate was obtained. Lysates were incubated on ice for 30 min with gentle mixing and then clarified by centrifugation at 17,000 × *g* for 10 min at 4L°C. The supernatant was transferred to a pre-chilled tube and diluted 1:1 with ice-cold dilution buffer (10 mM Tris-HCl pHL7.5, 150LmM NaCl, 0.5 mM EDTA). For immunoprecipitation, 25 µL of RFP-Trap Magnetic Agarose slurry per sample was equilibrated by washing once with 500 µL dilution buffer. Diluted lysates were added to the equilibrated beads and incubated for 1h at 4°C with gentle end-over-end rotation. Beads were then separated magnetically and washed three times with 500LµL wash buffer (10 mM Tris-HCl pHL7.5, 150 mM NaCl, 0.05% Nonidet P40 Substitute, 0.5LmM EDTA), transferring the beads to a fresh tube during the final wash to minimise nonspecific carryover. For mass spectrometry-based proteomic analysis, bound proteins were additionally washed (3×) with a detergent-free buffer (10 mM Tris, 150 mM NaCl, 0.5LmM EDTA, pH 7.5) followed by an overnight incubation at 37L°C in 100 mM HEPES buffer pH 8.0 supplemented with 5 mM Tris(2-carboxyethyl)phosphine, 10 mM 2-chloroacetamide, and 1 µg sequencing grade trypsin. Following on-bead digestion, peptide solutions were separated from the beads and acidified with TFA to a final concentration of 1%. For Western Blot analysis. bound proteins were eluted by resuspending beads in 80 µL 2× SDS sample buffer (120LmM Tris-HCl pHL6.8, 20% glycerol, 4% SDS, 0.04% bromophenol blue, 10% β-mercaptoethanol) and heating for 5 min at 95L°C. Eluted proteins were analysed by SDS–PAGE and Western blotting using an anti-mCherry antibody (Cat. ab167453; Abcam).

#### TMT-based proteomic identification of ALLO-1 binding partners

Following co-immunoprecipitation and on-bead digestion, the resulting tryptic peptides were labelled using an on-column TMT labelling protocol^48^. TMT-labelled samples were compiled into a single TMT sample and concentrated. Prior to LC-MS measurement, the peptide fractions were reconstituted in 0.1% TFA, 2% acetonitrile in water. Chromatographic separation was performed on an Easy-Spray Acclaim PepMap column 50Lcm longL×L75Lµm inner diameter (Thermo Fisher Scientific) at 55L°C by applying 120 min acetonitrile gradients in 0.1% aqueous formic acid at a flow rate of 300Lnl/min. An UltiMate 3000 nano-LC system was coupled to a Q Exactive HF-X mass spectrometer via an easy-spray source (all Thermo Fisher Scientific). The Q Exactive HF-X was operated in TMT mode with survey scans acquired at a resolution of 60,000 at m/z 200. Up to 18 of the most abundant isotope patterns with charges 2– 5 from the survey scan were selected with an isolation window of 0.7Lm/z and fragmented by higher-energy collision dissociation (HCD) with normalized collision energies of 32, while the dynamic exclusion was set to 35 s. The maximum ion injection times for the survey and MS/MS scans (acquired with a resolution of 30,000 at m/z 200) were 50 and 130 ms, respectively. The ion target value for MS was set to 3e6 and for MS/MS to 1e5, and the minimum AGC target was set to 1e3. The data were processed with MaxQuant v. 1.6.17.0^49^, and the peptides were identified from the MS/MS spectra searched against UniProt *C. elegans* reference proteome (UP000001940) using the built-in Andromeda search engine. Reporter ion MS2-based quantification was applied with reporter mass toleranceL=L0.003LDa and min. reporter PIFL=L0.75. Cysteine carbamidomethylation was set as a fixed modification, and methionine oxidation, glutamine/asparagine deamination, and protein N-terminal acetylation were set as variable modifications. For in silico digests of the reference proteome, cleavages of arginine or lysine followed by any amino acid were allowed (trypsin/P), and up to two missed cleavages were allowed. The FDR was set to 0.01 for peptides, proteins, and sites. Other parameters were used as pre-set in the software. Unique and razor peptides were used for quantification, enabling protein grouping (razor peptides are the peptides uniquely assigned to protein groups and not to individual proteins). Reporter intensity-corrected values were imported into Perseus v1.6.10.0^50^. Reverse hits, potential contaminants, and entries identified only by site were removed. Intensities were log₂-transformed, and only protein groups with complete quantitative data across all TMT channels were retained. Normalisation was performed by median subtraction within channels. PCA analysis performed on such processed data revealed that one of the control samples (ctrl_1) did not cluster well with the other two control samples (ctrl_2 and ctrl_3), plausibly due to an experimental error. The outlier control sample was excluded from analysis. ALLO-1 binding partners were defined as those meeting the threshold of a one-sided Student’s t-test (permutation-based FDR of 0.01 and S₀L=L0.5).

#### SDS-PAGE and Western Blotting

Protein samples were separated by SDS–PAGE on 10% polyacrylamide gels using a running buffer containing 25 mM Tris, 190 mM glycine, and 0.1% SDS. Electrophoresis was performed at a constant voltage of 120 V. Separated proteins were transferred onto PVDF membranes by wet transfer at 100 V for 1 h using a transfer buffer composed of 25 mM Tris, 190LmM glycine, and 10% methanol (pHL8.3). Total protein transfer was verified using the No-Stain Protein Labeling Reagent (Cat. A44717; Thermo Fisher Scientific) according to the manufacturer’s instructions. Membranes were then blocked for 45 min at room temperature in 5% (w/v) skimmed milk prepared in TBST buffer (50 mM Tris-HCl, 150 mM NaCl, 0.1% Tween-20, pHL7.5). Blots were incubated overnight at 4 °C with the primary anti-mCherry antibody (Cat. ab167453; Abcam) diluted in 5% milk/TBST. After three washes in TBST (10 min each), membranes were incubated for 1 h at room temperature with HRP-conjugated secondary antibodies diluted in the same blocking buffer. Protein bands were visualised using enhanced chemiluminescence on a ChemiDoc Imaging System (Bio-Rad).

#### Measurement of oxygen consumption rates (Seahorse XFp assay)

Oxygen consumption rates (OCR) were measured in *C. elegans* following a protocol adapted from^51^. Age-synchronised young adult wild-type and *allo-1* overexpression (OE) animals were collected from NGM plates seeded with *E. coli* OP50 and washed three times with M9 buffer by centrifugation (1,000 × *g*, 2 min) to remove bacteria. Approximately 10-15 worms were transferred into each well of an eight-well Seahorse XFp cell culture microplate (Agilent) pre-coated with Cell-Tak (Corning) to ensure adherence during measurement. Basal OCR was recorded using a Seahorse XFp Analyser (Agilent) with a standard measurement cycle consisting of 3 min mixing, 2 min waiting, and 2 min measurement. Mitochondrial uncoupling was induced by injection of 10 µM FCCP (carbonyl cyanide-4-(trifluoromethoxy)phenylhydrazone; Cat. SML2959, Sigma-Aldrich), and non-mitochondrial respiration was determined after subsequent injection of 40 mM sodium azide (Cat. S2002, Sigma-Aldrich). OCR values were normalised to the number of worms per well.

#### Worm motility assay

Age-synchronised young adult *C. elegans* wild-type and *unc-45(m94)* mutant animals were cultured at 15 °C and then shifted to 25 °C for 18-20 h to induce paralysis. Subsequently, worms were returned to 15 °C for approximately 10 h to allow partial motility recovery. For motility analysis, animals from each condition were transferred to unseeded NGM plates and spontaneous movements were recorded for 2 min using the WormLab imaging system (MBF Bioscience). Imaging parameters, including frame rate (7.5LframesLs⁻¹), exposure time, and gain, were kept constant across all recordings to ensure comparability. Movement trajectories were analysed in WormLab software (MBF Bioscience) to quantify motility parameters such as total track length and distance travelled per worm. Each assay included three independent biological replicates, with at least 75 animals analysed per replicate.

#### Fluorescence microscopy of polyQ40::YFP C. elegans

The number and average size of polyglutamine (polyQ) aggregates were quantified in *C. elegans* expressing *polyQ40::YFP* by imaging GFP fluorescence in young adult animals cultured at 20 °C. Fluorescence imaging was performed using an Axio Zoom.V16 stereomicroscope equipped with an Axiocam 705 monochrome CMOS camera (Carl Zeiss). Images were acquired in both brightfield and GFP channels under identical exposure settings. Normalised fluorescence intensity of whole animals was quantified using ImageJ (https://imagej.nih.gov/ij/). Data represent measurements from 30 worms per condition, pooled from three independent biological replicates.

#### Confocal microscopy

High-resolution imaging of *C. elegans* samples was performed using a ZEISS LSM800 laser scanning confocal microscope (Carl Zeiss Microscopy) equipped with 63×/1.4LNA oil-immersion and 10×/0.3LNA objectives. For all experiments, imaging parameters, including laser power, detector gain, and pixel size, were kept constant across samples to ensure quantitative comparability. Unless otherwise indicated, images correspond to single optical sections acquired under identical conditions. Image processing and analysis were carried out using ZEN (Zeiss) and ImageJ software. Only linear brightness and contrast adjustments were applied uniformly across the entire image for visualisation purposes without affecting the original data.

#### Image analysis

Image analysis was performed using ImageJ (https://imagej.nih.gov/ij/). Fluorescence intensity was quantified as mean grey value within defined regions of interest (ROIs). Mitochondrial morphology was analysed using the ImageJ Fiji plugin “Mitochondria Analyzer” v2.0.2 (https://sites.imagej.net/ACMito/). At least one ROI was analysed in a minimum of 20 images per condition across three biological replicates. For colocalization studies, the JaCoP plugin was applied^52^. Background subtraction was carried out using the rolling ball algorithm (radius = 50). Thresholds for the green and red channels were defined manually and maintained consistently across all images within a given experiment. Two ROIs containing individual cells per image were analyzed in 13-18 images per condition. All brightness and contrast adjustments were applied uniformly and exclusively for visualisation, without altering quantitative results or interpretation.

#### Transmission electron microscopy

Sample preparation and embedding for transmission electron microscopy (TEM) were performed as previously described^22^ with minor modifications. Briefly, C. elegans were immersed in 2.5% glutaraldehyde (EM grade, Cat. G7651, Sigma-Aldrich) prepared in 0.1 M PBS (pH 7.5). While in the fixative, worms were carefully nicked with a fine needle to facilitate better penetration of the fixative and then incubated overnight at 4 °C. After fixation, samples were rinsed three times with PBS and post-fixed on ice in a mixture of 1% aqueous osmium tetroxide (Polysciences Europe GmbH, 0972B-5) and 1.5% potassium ferrocyanide (Cat.

P3289, Sigma-Aldrich) for 30 min. The specimens were then treated with 1% aqueous thiocarbohydrazide (Cat. 88535, Sigma-Aldrich) for 40 min and post-fixed with 2% osmium tetroxide for 60 min at room temperature. Samples were subsequently incubated in 1% aqueous uranyl acetate (Cat. 6159-44-0, CHMES) at 4 °C overnight, followed by staining with 0.66% lead aspartate for 60 min at 60 °C. After dehydration through a graded ethanol series, samples were infiltrated with epoxy resin (Cat. 45-359, Sigma Aldrich), placed between two Aclar sheets (Cat. L4458, Agar Scientific) or positioned into flat silicone molds to enable further animals’ cross sections. Polymerisation was done at 60 °C for 72 h. Once the resin blocks were cured and the Aclar removed, samples were trimmed and sectioned using an ultramicrotome (EM UC7, Leica). Ultrathin sections (70 nm) were collected on 200-mesh nickel grids (Cat. G2200N, Agar Scientific) or copper slot grids (Cat. FCF 2010-CU-SB-50, Electron Microscopy Sciences). Sections were imaged using a Tecnai T12 BioTwin transmission electron microscope (FEI, Hillsboro, OR, USA) equipped with a 16-megapixel TemCam-F416(R) camera (TVIPS GmbH).

## ACKNOWLEDGMENTS

We thank the Genome Engineering Unit and the Microscopy and Cytometry Facility at the International Institute of Molecular and Cell Biology in Warsaw (IIMCB) for their technical support. We are especially grateful to Dr. Matylda Macias and Aleksandra Szybińska for assistance with transmission electron microscopy. Access to core facilities was provided through the IIMCB IN-MOL-CELL Infrastructure, funded by the European Union’s NextGenerationEU programme under the National Recovery and Resilience Plan. IN-MOL-CELL also received support from the European Union through Horizon Europe (Project 101059801, RACE) and from the RACE-PRIME project implemented within the IRAP programme of the Foundation for Polish Science, co-financed by the European Funds for Smart Economy 2021-2027 (FENG). Proteomic measurements were performed at the Proteomics Core Facility, IMol Polish Academy of Sciences. We extend our gratitude to Dr. Dorota Stadnik, Dr. Magdalena Chojnacka, and Mrs Viktoriia Lastivka for assistance with sample preparation and LC-MS/MS measurements. We thank Prof. Johannes Buchner (Technical University of Munich, Germany) for providing the anti-SIP-1 antibodies and Prof. Nektarios Tavernarakis (Institute of Molecular Biology and Biotechnology, Foundation for Research and Technology-Hellas, Crete, Greece) for providing the *C. elegans* mitoRosella strain. Seahorse XFp instrument access and training were kindly provided by Altium International. We also thank Marta Niklewicz for her exceptional assistance with *C. elegans* culture maintenance and preparation of nematode growth media. Finally, we thank all members of the Pokrzywa laboratory for helpful discussions, experimental advice, and critical feedback on the manuscript.

## Declaration of generative AI and AI-assisted technologies in the writing process

During the preparation of this work the authors used ChatGPT to improve the readability and language of the manuscript. After using this tool, the authors reviewed and edited the content as needed and take full responsibility for the content of the published article.

## AUTHOR CONTRIBUTIONS

**A.S.:** Conceptualization; Data curation; Formal analysis; Methodology; Investigation; Validation; Visualization; Writing-review & editing. **L.B.:** Formal analysis; Methodology; Investigation; Visualization. **N.A.S.**: Formal analysis; Visualization; Writing-review & editing. **P.T.**: Validation. **R.S.:** Formal analysis. **W.P.:** Conceptualization; Formal analysis; Funding acquisition; Project administration; Resources; Supervision; Writing-original draft.

## Supplemental figures legends

**Figure S1.**
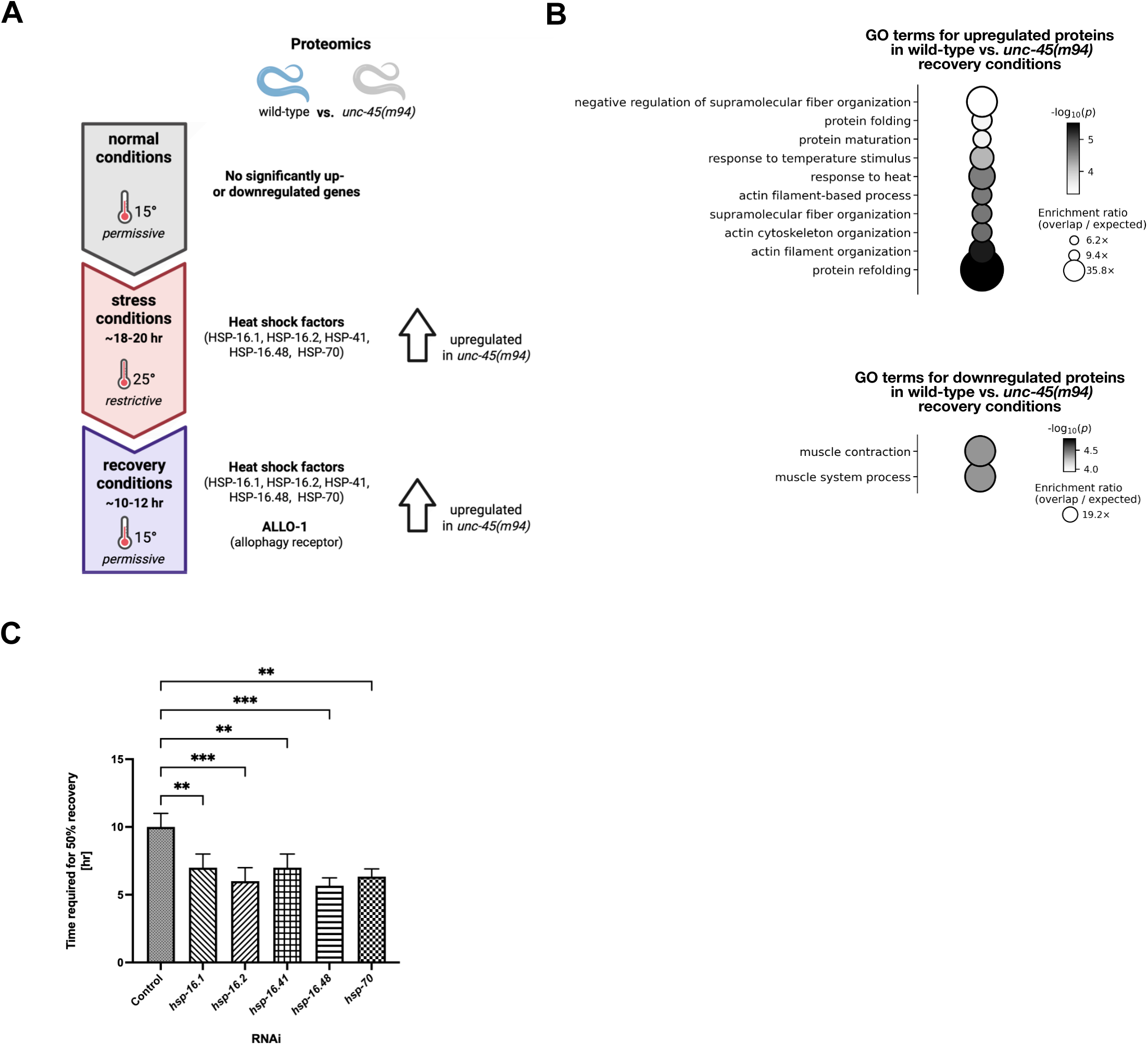
Proteomic and genetic analyses of recovery dynamics in *unc-45(m94)* mutants. **(A)** Schematic summary of the proteomics workflow comparing young adult wild-type and *unc-45(m94)* animals maintained at 15 °C, shifted to 25 °C, and after recovery at 15 °C. **(B)** GO enrichment analysis of biological process terms for proteins up- or downregulated in *unc-45(m94)* animals during recovery. Up - and downregulated proteins were defined as those showing ≥ 0.5-fold or ≤ −0.5 enrichment, respectively, relative to wild-type in addition to meeting the initial permutation-based *t*-test criteria (two-sided, S₀ = 0.1, FDR = 0.05). GO analysis was performed using WebGestalt (FDR = 0.1, Benjamini-Hochberg method). Circle size denotes enrichment ratio; colour shading indicates statistical significance (-log_10_ p). **(C)** Quantification of recovery time (hours required for ∼50 % of the population to regain motility) for young adult *unc-45(m94)* animals after down-shift from 25 °C to 15 °C under control (empty vector), *hsp-16.1*, *hsp-16.2*, *hsp-16.41*, *hsp-16.48*, *hsp-70*, or *allo-1* RNAi. RNAi was initiated at the early L4 stage. Data represent mean ± SEM from ≥ 3 biological replicates. Statistical significance was assessed using one-way ANOVA with Dunnett’s multiple-comparisons test (* PL<L0.05, ** PL<L0.01, *** PL<L0.001, **** PL<L0.0001).

**Figure S2.**
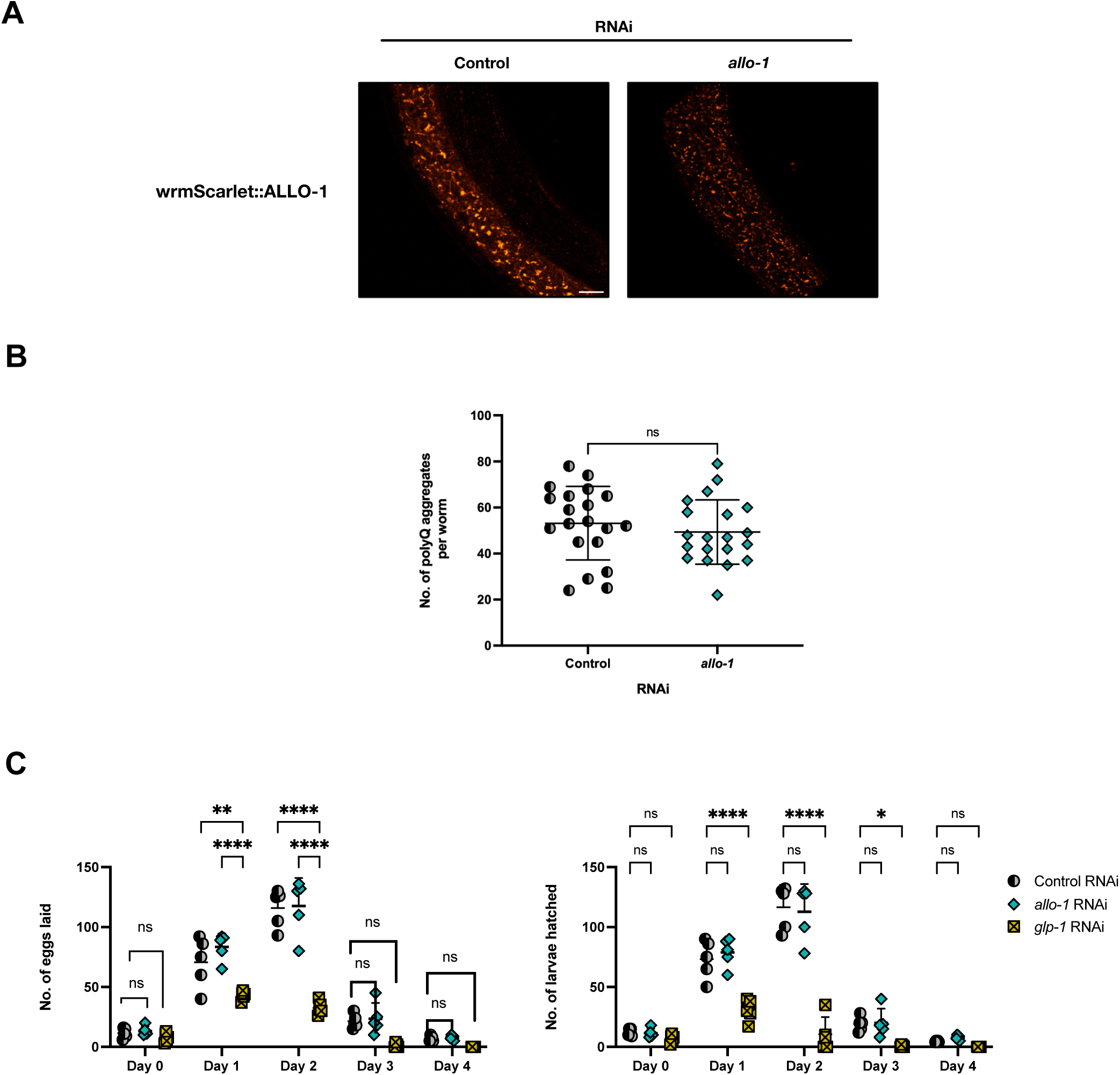
Control experiments validating ALLO-1 reporter specificity and excluding indirect effects. **(A)** Validation of *allo-1* RNAi specificity. Representative confocal images of wrmScarlet::ALLO-1 animals subjected to control or *allo-1* RNAi initiated at the early L4 stage, at 20 °C. Scale bar, 10 µm. **(B)** Quantitative analysis of proteostasis capacity in body wall muscle using the polyQ40::YFP aggregation reporter. Shown are the total number of fluorescent aggregates per animal under control or *allo-1* RNAi conditions initiated at early L4, at 20 °C. Data represent ≥ 30 animals from three biological replicates. Statistical significance was assessed by the Mann-Whitney test (ns - not significant). **(C)** Assessment of fertility following *allo-1* or *glp-1* RNAi. Left - total number of eggs laid per day by wild-type animals (two animals per plate) under control, *glp-1*, or *allo-1* RNAi initiated at the early L4 (n = 30 from three independent biological replicates). Right - total number of larvae hatched from the same plates, at 20 °C. Data represent mean ± SEM. Statistical analysis used two-way ANOVA with Dunnett’s or Tukey’s multiple-comparisons test as indicated (ns - not significant; * *P* < 0.05; ** *P* < 0.01; *** *P* < 0.001 **** *P* < 0.0001).

**Figure S3.**
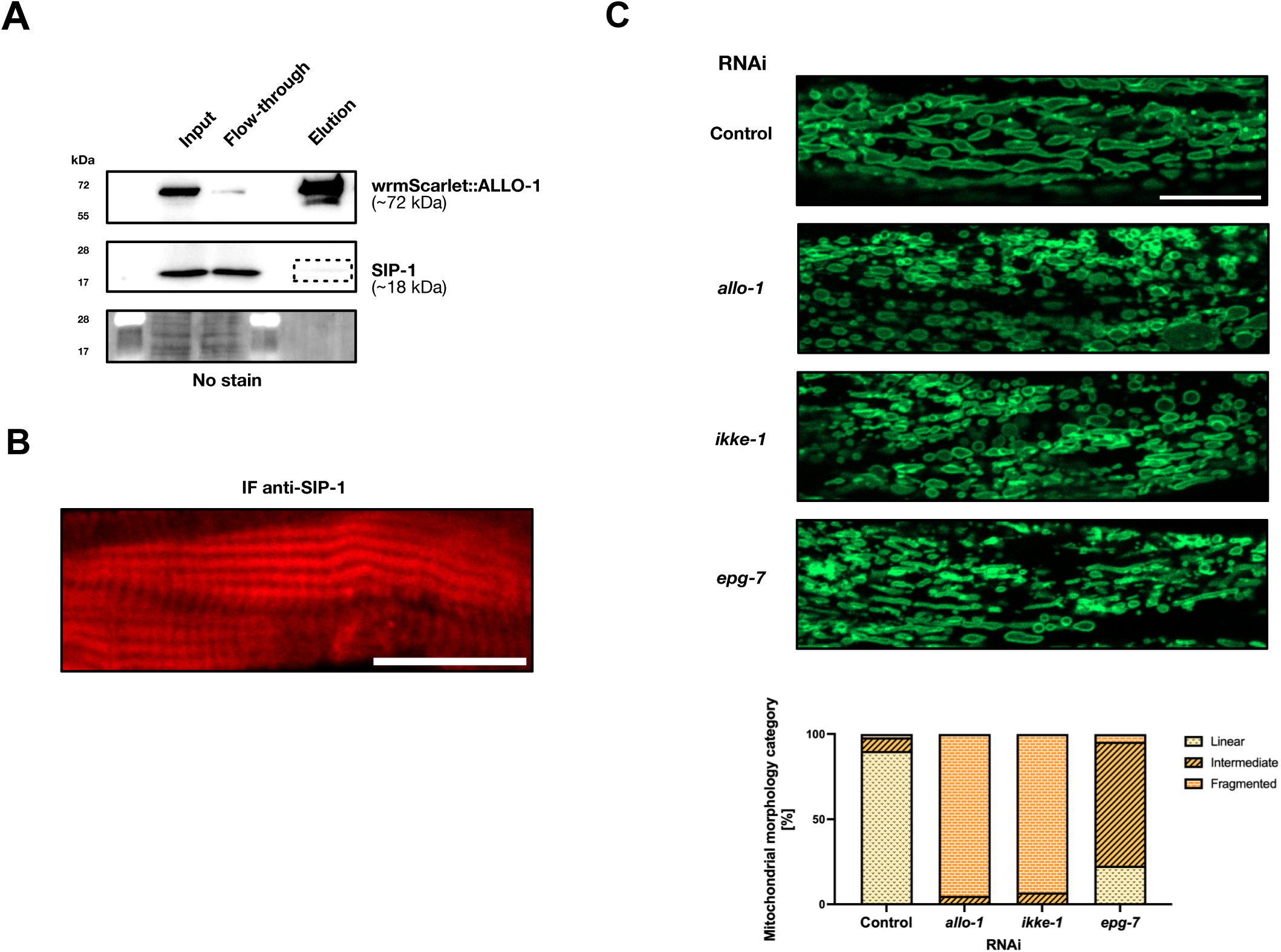
Validation of ALLO-1 interactors and analysis of mitochondrial morphology. **(A)** Western blot showing detection of wrmScarlet::ALLO-1 and SIP-1 in immunoprecipitation eluates using anti-mCherry and anti-SIP-1 antibodies. Equal protein loading was confirmed using No-Stain Protein Labeling Reagent. **(B)** Representative confocal image showing SIP-1 localization in body wall muscle of young adult wild-type animals stained with anti-SIP-1 antibody. Scale bar, 10Lµm. **(C)** Top - representative confocal images of body wall muscle mitochondria in young adult wild-type animals expressing TOMM-20::GFP under control, *allo-1*, *ikke-1*, and *epg-7* RNAi, initiated at the early L4 stage, at 20 °C. Scale bar, 10Lµm. Bottom - quantification of animals displaying linear, intermediate, or fragmented mitochondrial morphology (n = 20 images per condition per biological replicate; three biological replicates).

**Figure S4.**
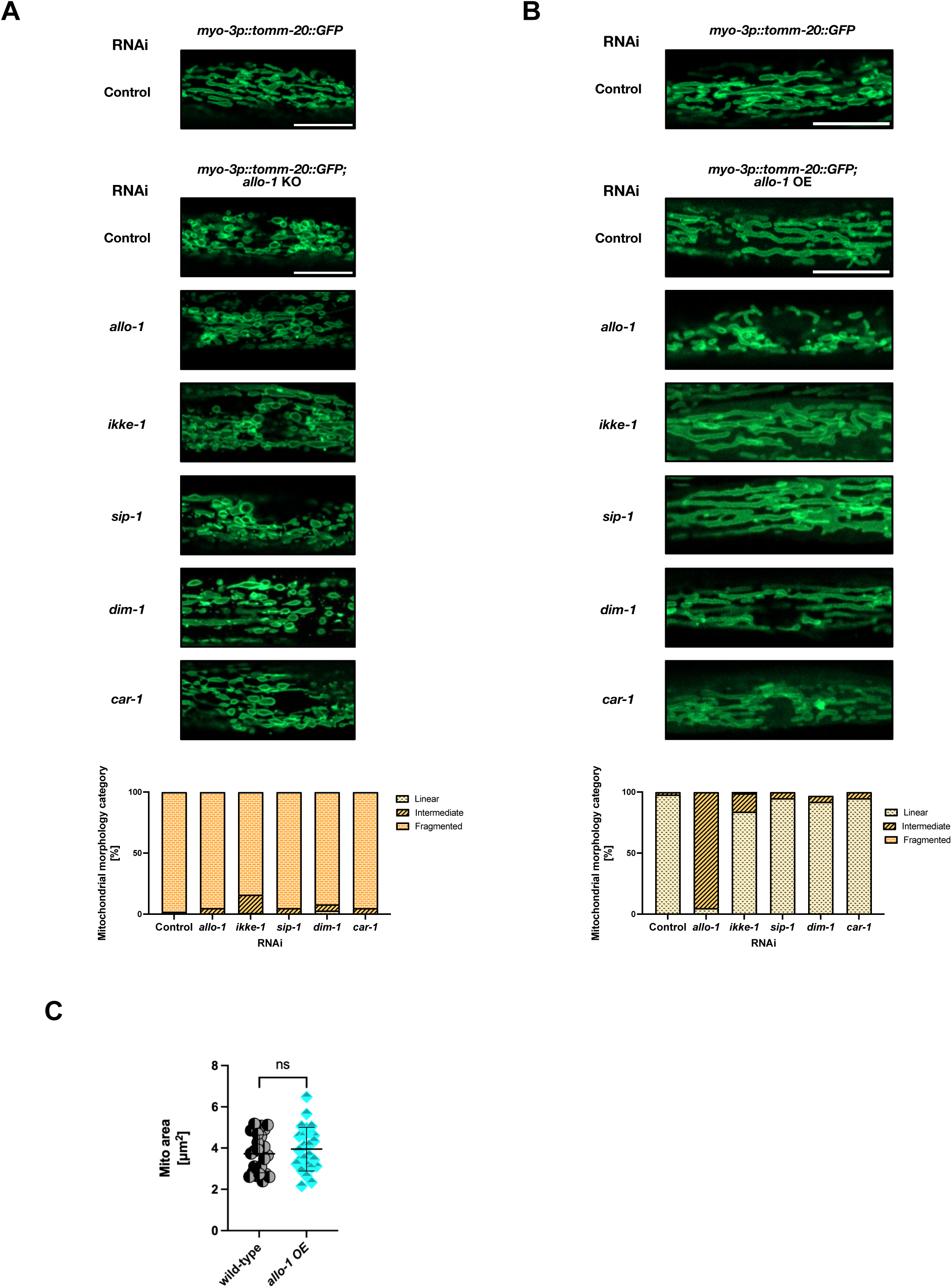
Mitochondrial morphology in ALLO-1 knockout and overexpression strains following RNAi of ALLO-1 network components. **(A)** Top - representative confocal images of body wall muscle mitochondria in young adult *allo-1* knockout (KO) animals expressing TOMM-20::GFP, following control, *allo-1*, *ikke-1*, *sip-1*, *dim-1*, or *car-1* RNAi, initiated at the early L4 stage, at 20 °C. Scale bar, 10 µm. Bottom - quantification of the proportion of animals displaying linear, intermediate, or fragmented mitochondrial morphology (n = 20 images per condition per biological repeat; three biological repeats). **(B)** Top - representative confocal images of body wall muscle mitochondria in young adult *allo-1*-overexpression (OE) animals expressing TOMM-20::GFP, following control, *allo-1*, *ikke-1*, *sip-1*, *dim-1*, and *car-1* RNAi, initiated at the early L4 stage, at 20 °C. Scale bar, 10 µm. Bottom - quantification of animals displaying linear, intermediate, or fragmented mitochondrial morphology (n = 20 images per condition per biological replicate; three biological replicates). **(C)** Quantitative analysis of mitochondrial area in body wall muscle of young adult wild-type and *allo-1* OE animals expressing TOMM-20::GFP initiated at L4, at 20 °C. Mito=mitochondria. Data represent mean ± SEM (n ≥ 3 biological replicates). Statistical significance was assessed by Welch’s t-test. (ns - not significant).

